# Lytic bacteriophages interact with respiratory epithelial cells and induce the secretion of antiviral and proinflammatory cytokines

**DOI:** 10.1101/2024.02.06.579115

**Authors:** Paula F. Zamora, Thomas G. Reidy, Catherine R. Armbruster, Ming Sun, Daria Van Tyne, Paul E. Turner, Jonathan L. Koff, Jennifer M. Bomberger

## Abstract

Phage therapy is a therapeutic approach to treat multidrug resistant infections that employs lytic bacteriophages (phages) to eliminate bacteria. Despite the abundant evidence for its success as an antimicrobial in Eastern Europe, there is scarce data regarding its effects on the human host. Here, we aimed to understand how lytic phages interact with cells of the airway epithelium, the tissue site that is colonized by bacterial biofilms in numerous chronic respiratory disorders. We determined that interactions between phages and epithelial cells depend on specific phage properties as well as physiochemical features of the microenvironment. Although poor at internalizing phages, the airway epithelium responds to phage exposure by changing its transcriptional profile and secreting antiviral and proinflammatory cytokines that correlate with specific phage families. Overall, our findings indicate that mammalian responses to phages are heterogenous and could potentially alter the way that respiratory local defenses aid in bacterial clearance during phage therapy. Thus, besides phage receptor specificity in a particular bacterial isolate, the criteria to select lytic phages for therapy should be expanded to include mammalian cell responses.

## INTRODUCTION

Phage therapy is a re-emerging therapeutic alternative to treat multidrug resistant (MDR) infections that relies on the use of bacteriophages (or phages) that infect and kill specific bacterial strains (1). Even though phage therapy is increasingly being used in clinical trials and compassionate use cases (via Emergency Use Authorizations) in the United States (2), there are decades of experience with phage therapy in Georgia, Poland, and Russia (3), where it is routine medical practice to prescribe phages for difficult-to-treat infections (4). Thanks to the success of phage therapy in Western Europe and the research from the Eliava Institute of Bacteriophages, Microbiology and Virology (Tblilisi, Georgia) and the Ludwik Hirszfeld Institute of Immunology and Experimental Therapy (Wroclaw, Poland), there has been a renewed interest in this therapeutic strategy in the Western world. Their expertise in phage isolation, preparation, and application has been crucial for the planning of clinical trials and expanding the access to phage therapy (5). Considering the problem of increasing bacterial antibiotic resistance (6) and the decrease of new of compounds in the antibiotic pipeline (7), it is imperative that we explore phage therapy as a potential treatment for infections in the United States.

Some of the excitement towards phage therapy comes from it being considered relatively safe, with few side effects reported (8). Phage therapy preferentially uses lytic phages, and it is thought to be innocuous in the human host because phages are unable to replicate inside mammalian cells (1, 9). While true, prior reports have indicated that bacteriophages can bind and be taken up by eukaryotic cells (10–12). This knowledge raises concerns regarding phages in a therapeutic context, as high doses of phages are often needed to elicit a therapeutic effect (1, 13–15). Moreover, because of their self-replicating ability, phage concentrations increase during treatment until the number of susceptible bacteria has decreased to a point that phage replication can no longer be supported. As a result, mammalian mucosal surfaces are likely in contact with phages during therapy.

Even though phage therapy is being used to treat various MDR infections (2), one of its most promising applications is towards the treatment of *Pseudomonas aeruginosa*. *P. aeruginosa* is a bacterium cataloged by the Center for Disease Control and Prevention (CDC) to be a serious threat to human health because MDR strains have emerged that are resistant to almost all available antibiotics (16). As an opportunistic pathogen, *P. aeruginosa* infections are particularly risky in vulnerable populations, such as hospitalized patients or people with weakened immune systems (17). People with cystic fibrosis (pwCF), a genetic disease characterized by the buildup of thick and sticky mucus in the lungs, commonly harbor lifelong chronic *P. aeruginosa* infections that require recurrent antibiotic treatment (18). Infections with *P. aeruginosa* correlate with declines in lung function in pwCF and are the leading cause of respiratory failure and mortality in this population (19). For these reasons, there is a critical need to find therapeutic alternatives to treat *P. aeruginosa* infections, and phage therapy is such an option.

Currently, treatment of lung infections is one of the most common indicators for phage therapy (20). However, key gaps in knowledge related to the interactions between phages and the lung need to be answered before phage therapy becomes more prevalent. Information regarding the immunogenicity of lytic phages is needed to be able to make better phage choices during therapy (20). In addition to plasma membrane receptor-ligand specificity, mammalian cells uptake viruses and other nanoparticles in a size- and shape-dependent manner (21, 22). Phages of identical bacterial specificity can have different physical features, and thus, interact with the human host in a diverse fashion. To better understand how the lung responds to phage therapy, we examined associations between a panel of lytic bacteriophages with therapeutic potential and respiratory epithelial cells. We showed that phages interact with bronchial epithelial cells in a manner that depends on intrinsic morphological phage features, in addition to specific properties of the mucosal microenvironment. Bronchial epithelial cells uptake phages at slower rates than other reported cell types (11), and some phages can cross to the basolateral space, suggesting that selected phages could enter the bloodstream during inhaled phage therapy. Even though uptake is rare, the airway epithelium detects phages and mounts a transcriptomic response that is enriched in the expression of immune-associated genes. Together, our findings indicate that respiratory epithelial cells sense and respond to lytic phages. Consequently, phage therapy could induce changes in the airways through a mechanism that is independent of the direct effect of phages in reducing bacterial loads. We propose that these changes could be anticipated before phage administration to potentiate the effect of this therapeutic.

## RESULTS

### Panel of therapeutic *P. aeruginosa* lytic bacteriophages is genetically and morphologically diverse

To model phage therapy in the CF airways, we utilized a panel of *P. aeruginosa* phages that have been used therapeutically in pwCF under FDA emergency use authorization and in a clinical trial (NCT04684641) to treat chronic MDR *P. aeruginosa* infections (23). We complemented this panel with additional *P. aeruginosa* phages obtained from hospital wastewater that have lytic activity against various *P. aeruginosa* clinical isolates from pwCF that were isolated for use in therapeutic applications (24). After initial characterization for their ability to replicate at high titers in large volume *P. aeruginosa* cultures, we chose a panel of four phages for further studies. These phages are named OMKO1, LPS-5, PSA04, and PSA34 (Fig. S1). To obtain highly pure phage particles to utilize in mammalian cell culture studies, we developed a protocol based on the use of several filtration steps, nuclease treatments, and two consecutive cesium chloride (CsCl) gradient centrifugations that lead to phage preparations mostly devoid of bacterial proteins, bacterial nucleic acids, and endotoxins (Table S1). Notably, the CsCl gradients enabled separation of phage in the density gradient from contaminating bacterial products, based on their density differences. Additionally, the phage bands were collected by puncturing the ultrathin wall of the centrifugation tubes, preventing cross-contamination between phages and bacterial debris (Fig. S1).

To confirm their lytic activity, we conducted killing assays using a panel of laboratory-adapted strains and clinical isolates of *P. aeruginosa* grown in abiotic planktonic and biofilm conditions, as well as biofilms grown in association with CF human airway epithelial cells (AECs, CFBE41o-cell line, a homozygous ΔF508 CFTR bronchial epithelial cell line) (Fig. 1A, Fig. 1B, Fig. S2, and Table S2) (25). Overall, we found that LPS-5 and PSA34 had better killing activity than OMKO1 and PSA04, although the degree of killing for each phage-bacterium pair was different for each assay. To determine the morphology and structural features of these phages, we conducted transmission electron microscopy (TEM) of purified phage stocks (Fig. 1C and Fig. 1D). Phages OMKO1 and LPS-5 were found to be myoviruses, whereas PSA04 and PSA34 were established as a siphovirus and a podovirus, respectively. The phages in our panel had different dimensions, with OMKO1 being a jumbo phage of 400 nm in length in its longest axis that is much larger in size than the other phages (Fig. 1D). Phages LPS-5, PSA04, and PSA34 measured 150, 200, and 80 nm in their longest axis, respectively. While their head sizes were similar, they differed in their tail properties. LPS-5 had a rigid tail, whereas PSA04 had a longer and more flexible tail, while PSA34 had a very short tail. Given that structural features of phage particles are key to defining their interactions with the mammalian cells they encounter (11), these data suggest that the different phages in our panel might elicit different mammalian cell responses.

**Figure 1.**
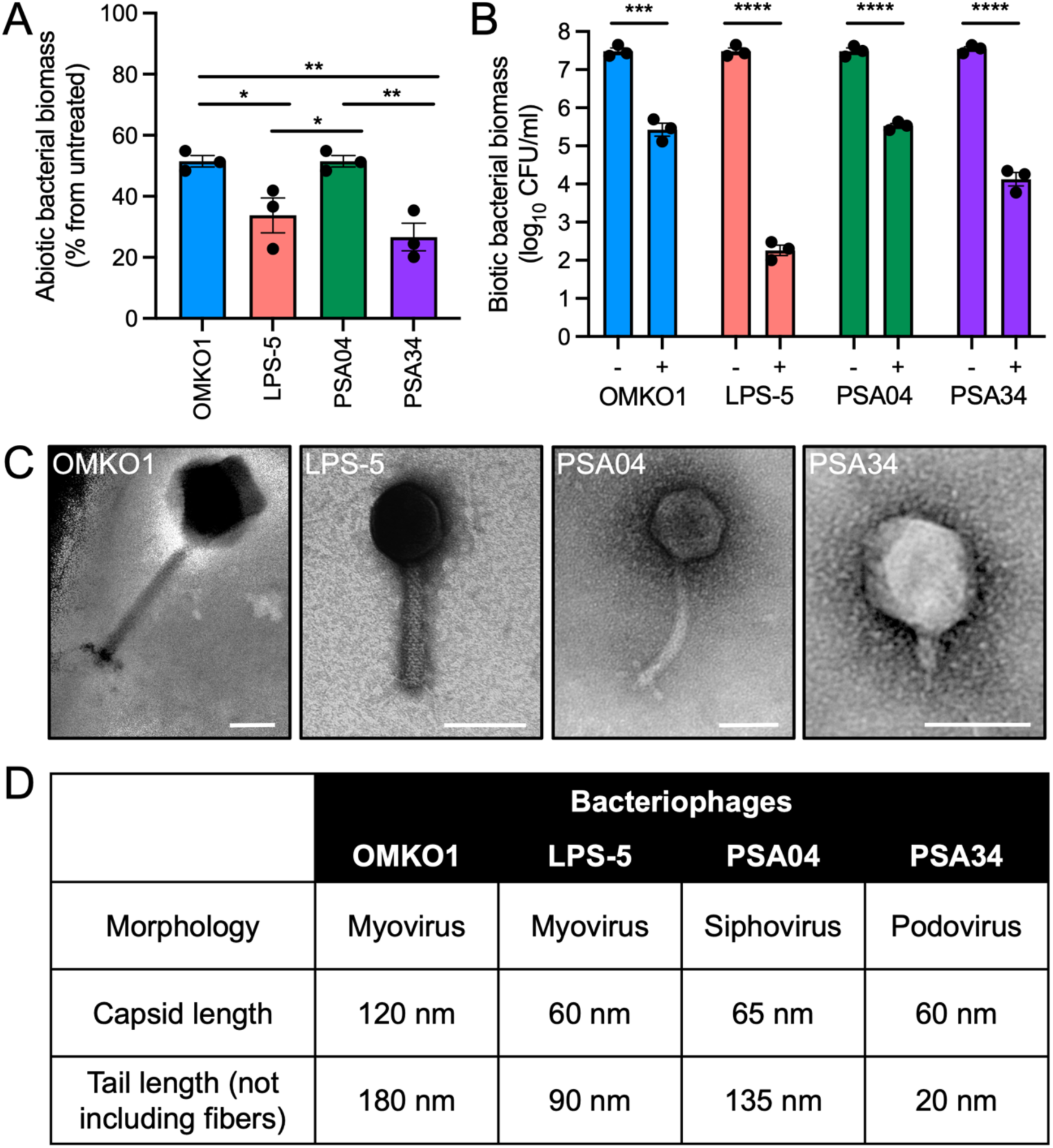
Phage panel represents lytic phages of different morphologies with clinical relevance. (A) *P. aeruginosa* biofilms for strains PAO1, DVT411, and DVT423 were grown on plastic wells in LB medium for 18 h. Phages were added at a concentration of 0.1 PFU/cell for 6 h. Results are shown as percentage compared to phage-untreated wells. (B) PAO1, DVT411, and DVT423 were grown on CFBE41o-cells for 16 h in MEM. Cells were apically treated with phages OMKO1, LPS-5, PSA04, and PSA34 at a concentration of 1 PFU/cell for 8 h. Bacterial aggregates were solubilized, and bacterial cells were separated from phages by centrifugation. Biomass was quantified by CFU drip assays. (C) Representative images of transmission electron microscopy pictures of negatively stained phages. Scale bar, 50 nm. (D) Table depicting phage structural characteristics. Phages were classified according to their morphology. Capsid and tail length measurements were obtained using ImageJ using the average lengths from three independent images. Error bars indicate SEM for three independent biological replicates. *p<0.05, **p<0.01, ***p<0.001, ****p<0.000.1.

To evaluate phage taxonomy and genomic features, we conducted whole genome sequencing of phage DNA, assembled and annotated genomes, and compared them to one another and to previously sequenced phages using BlastN (Table S1). We determined the phylogenetic relationship between our panel of phages and other lytic phages by conducting a literature search of phages of known morphology and with available genome sequences that also have a preferred bacterial host whose genome sequence was available in NCBI at the time of this manuscript submission (Fig. 2). Phages OMKO1, LPS-5, and PSA34 clustered with phages of similar morphology and genome length. In contrast, phage PSA04 was more genetically divergent to the other phages than the rest of our phage panel and did not cluster with any of the other siphoviruses analyzed. Interestingly, within clusters of phages of similar morphology and genome length, head and tail lengths varied, suggesting that phylogenetically related phages are not necessarily morphologically similar. Taken together, the phages in this panel have different lytic characteristics, morphology, and genome features, representing the variety of phages used clinically to treat chronic *P. aeruginosa* infections.

**Figure 2.**
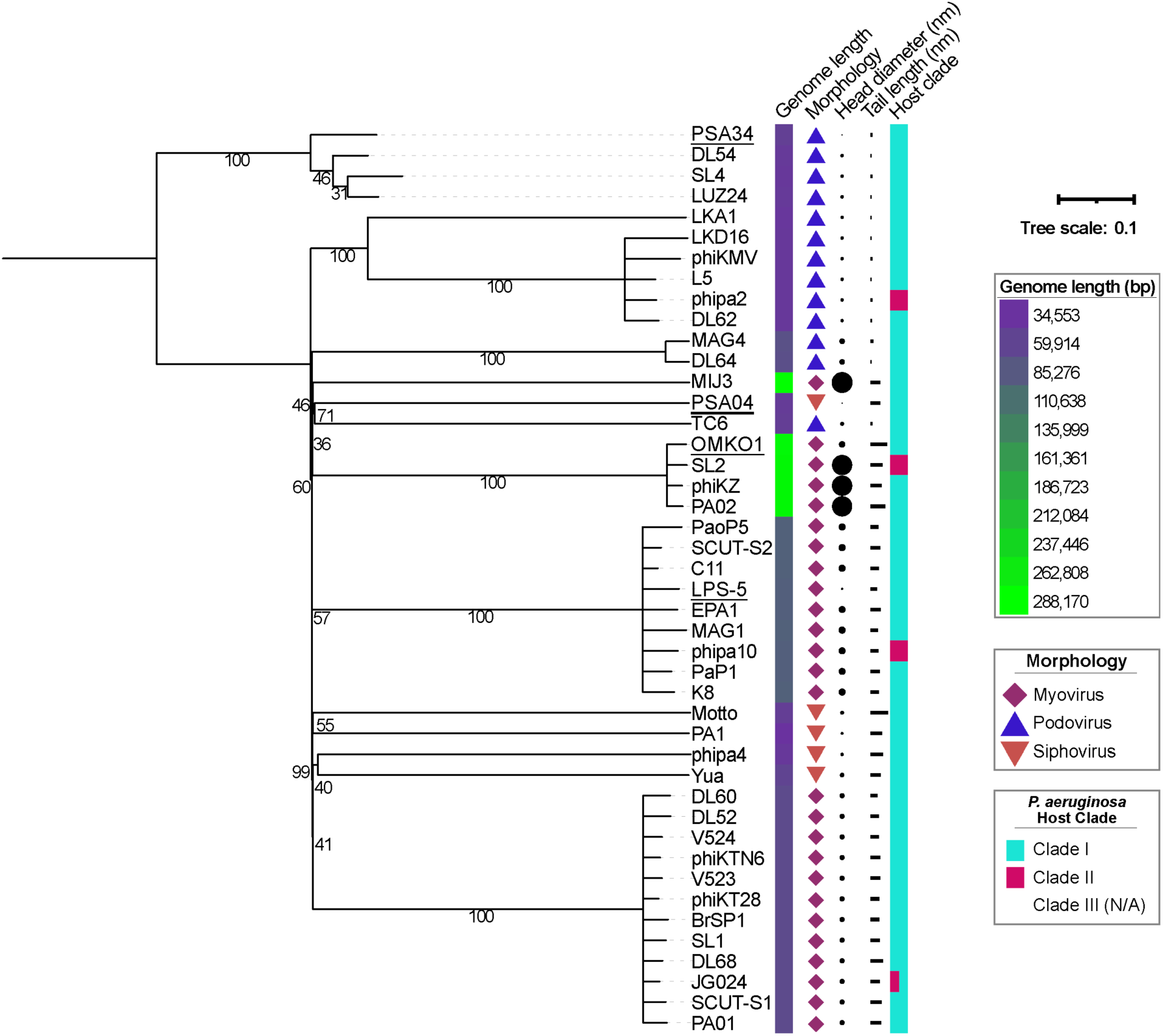
Panel of phages are phylogenomically and morphologically distinct. Midpoint rooted tree depicting phylogenomic distances between phage, with the numbers on branches indicating the bootstrap values. Branches with bootstrap values < 50 were collapsed. Phages were chosen if their genome sequences and the ones of their host were accessible, and if TEM images were available. Head and tail lengths on their longest axis were measured using ImageJ. The scale bar represents the number of amino acid substitutions per site. Phages used in this study are underlined.

### Phage interactions with the airway epithelium are favored under acidic pH and in areas of epithelial remodeling

We next sought to evaluate the ability of phages to associate with the airway epithelium. We tested whether acidic pH influenced phage interactions with bronchial epithelial cells, as the acidic pH of the airway surface liquid (ASL) is a characteristic of the pathophysiology of CF and other inflammatory disorders, such as chronic obstructive pulmonary disease (COPD) (26–29). Phages OMKO1, LPS-5, PSA04, and PSA34 were incubated apically with CF AECs for 24 hours (h) in cell culture media that was pH-adjusted to 6.5 or 7.5, and phage association with the airway epithelium was determined by quantifying remaining viable phage after washing by plaque assay (Fig. 3A and Fig. S3). We determined that pH 6.5 promoted phage interactions with mammalian cells to a greater extent than pH 7.5. The increase in cell-association was statistically significant for the myoviruses OMKO1 and LPS-5, suggesting that virion morphology is a factor contributing to the pH-dependency in epithelial-phage interactions. We complemented these studies by conducting confocal microscopy imaging of fluorescently-labeled phages after treating CF AECs at pH 6.5 or 7.5 for 24 h (Fig. 3B, Fig. 3C, and Fig. S3). We observed that acidic pH promoted particle aggregation on the airway epithelium. We quantified the total number of particles associated with respiratory cells and determined that for phages LPS-5 and PSA34, acidic pH tended to promote a higher number of associated particles compared to incubation at pH 7.5, and this increased association was statistically significant for phage OMKO1. pH did not affect the total number of associated particles for phage PSA04, which in general, poorly associated with mammalian cells compared to the other phages in the panel. When binning for aggregate size, we found that the proportion of particles of smaller sizes was higher at pH 7.5 compared to pH 6.5, suggesting that the local pH in mucosal surfaces influences phage-phage interactions (Fig. 3D and 3E). Phages PSA04 and PSA34 had a significantly higher proportion of particles over 5 µm^3^ at pH 6.5 compared to OMKO1 and LPS-5, suggesting that myoviruses are less prone to aggregation on epithelial cells. Importantly, phage particle dynamics in the absence of the airway epithelium is comparable between both pH values, suggesting that phage aggregation depends on the airway mucosal surface (Fig. S4). Overall, our experiments indicate that phage associations with the airway epithelium are pH dependent and specific for different phage families.

**Figure 3.**
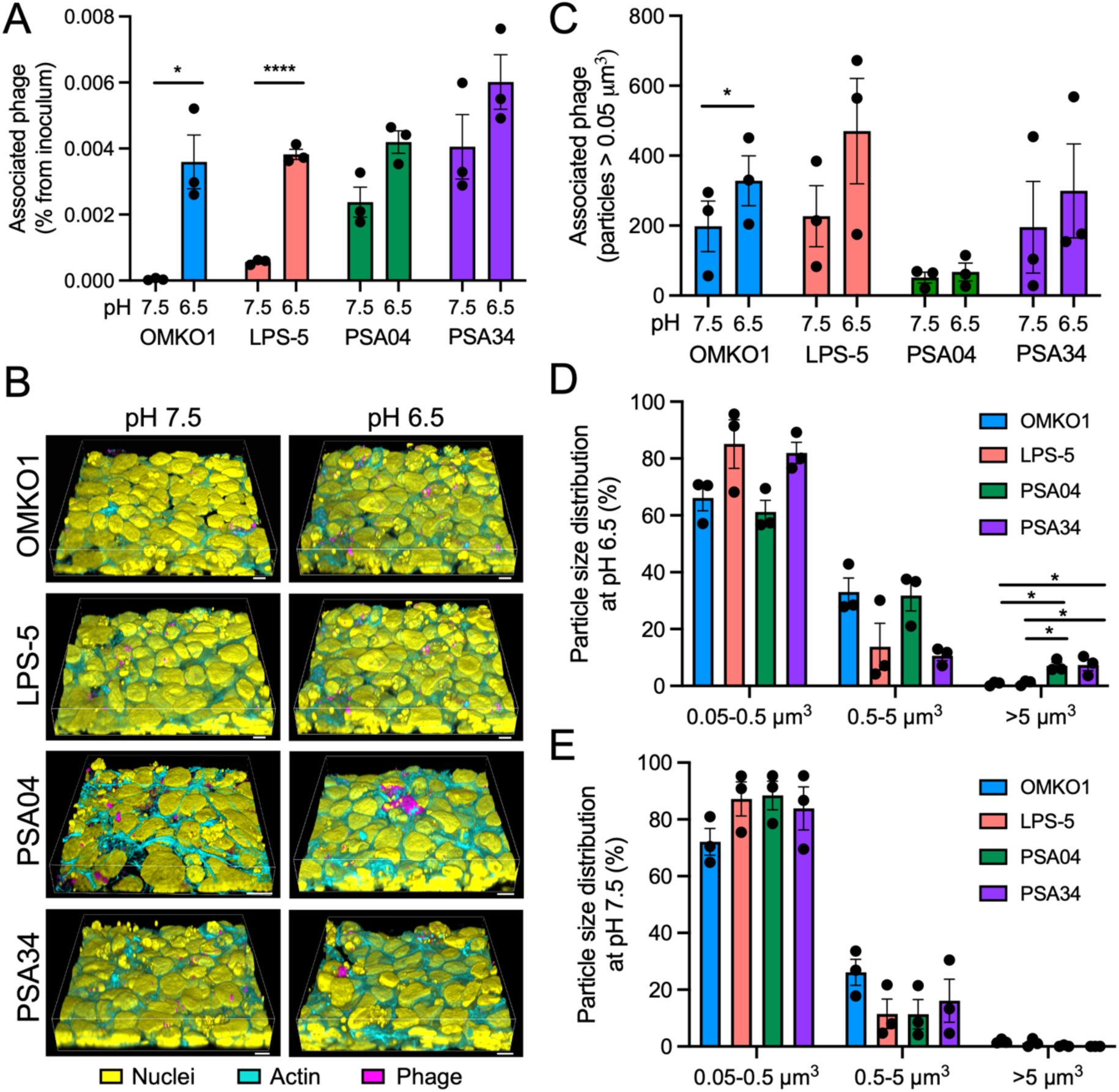
Phage interactions with the airway epithelium are pH dependent. (A) CFBE41o-cells were apically treated with phages OMKO1, LPS-5, PSA04, and PSA34 at a concentration of 1 x 10^9^ PFU/ml in MEM pH-adjusted to 6.5 or 7.5 for 24 h. Cells were washed extensively, and phage viable particles were quantified by plaque assay using the double-agar method. (B to D) CFBE41o-cells were apically treated with fluorescently labeled phages OMKO1, LPS-5, PSA04, and PSA34 (magenta) at a concentration of 1 x 10^9^ PFU/ml in MEM pH-adjusted to 6.5 or 7.5 for 24 h. Cells were washed and fixed for confocal microscopy. Cells were stained with fluorescently labeled phalloidin (cyan) and Hoechst (yellow) to label actin and nuclei, respectively. (B) 3D reconstructions of confocal microscopy Z-stacks. Scale bar, 10 µm. (C to D) TRITC-positive particles were identified in 3D reconstructions using the 3D Object Measurement function in Nikon Elements. (C) Total number of particles larger than 0.05 µm^3^ for both pH conditions. (D and E) Particles over 0.05 µm^3^ were binned in three groups: 0.05-0.5 µm^3^, 0.5-.5 µm^3^, and over 5 µm^3^. Error bars indicate SEM for three independent biological replicates. *p<0.05, ****p<0.0001.

When imaging phage-epithelial interactions, we observed that phages were repeatedly found in areas rich in condensed nuclear material. These manifestations in epithelial tissues are often associated with cellular remodeling due to cell extrusion, a homeostatic tissue process where cytoskeletal proteins contract around apoptotic or overcrowded cells and remove them from the epithelial layer without disrupting barrier integrity (30, 31). To better understand whether phages localize more frequently in distinct epithelial regions, we imaged CF AECs after a 1 h apical treatment with our panel of phages, a time chosen to exclude the possibility of phages inducing cellular remodeling, as the kinetics of epithelial extrusion is slower than that time frame (Fig. 4A). We found that all phages in our panel associated with the epithelium in areas of increased actin and DNA staining, suggesting that phages might bind cellular components that are exposed or released during epithelial extrusion. In imaging experiments where we stained for phosphatidylserine, a plasma membrane phospholipid that is flipped to the outer leaflet during apoptosis, we found that phages associated in areas that were both positive and negative for phosphatidylserine staining (Fig. S5). This observation suggests that phosphatidylserine is not required for phage-epithelial association.

**Figure 4.**
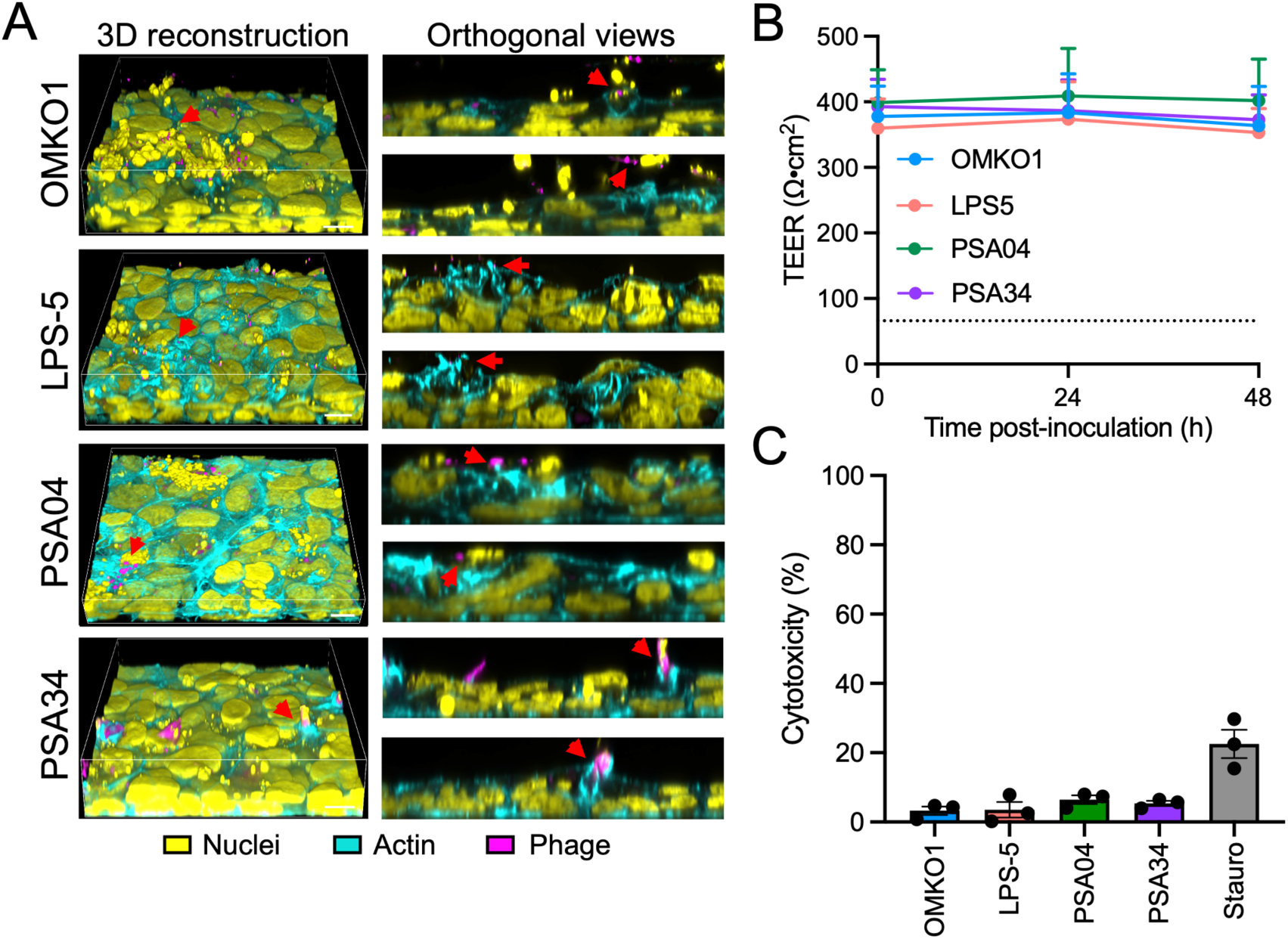
Phages locate to areas of epithelial remodeling without disrupting cellular integrity. (A) CFBE41o-cells were apically treated with fluorescently labeled phages OMKO1, LPS-5, PSA04, and PSA34 (magenta) at a concentration of 1 x 10^9^ PFU/ml in MEM pH-adjusted to 6.5 for 1 h. Cells were washed and fixed for confocal microscopy. Cells were stained with fluorescently labeled phalloidin (cyan) and Hoechst (yellow) to label actin and nuclei, respectively. Images are shown as 3D reconstructions of confocal microscopy Z-stacks (left), XZ and YZ projections (right). Red arrowheads indicate areas rich in actin staining and nuclear extracellular material. Scale bar, 10 µm. (B) CFBE41o-cells were apically treated with phages OMKO1, LPS-5, PSA04, and PSA34 at a concentration of 1 x 10^9^ PFU/ml in MEM pH-adjusted to 6.5 for 48 h. At 0, 24, and 48 h post-treatment, transepithelial electrical resistance (TEER) was measured. Dotted line indicates the TEER value for cells with disrupted integrity. Values are shown as the read value for each treatment minus the value of a Transwell insert without cells. (C) CFBE41o-cells were apically treated with phages OMKO1, LPS-5, PSA04, and PSA34 at a concentration of 1 x 10^9^ PFU/ml in MEM pH-adjusted to 6.5 for 24 h. Supernatants were collected and glucose-6-phosphate dehydrogenase activity was quantified by measuring the reduction of resazurin to resorufin by fluorescence. Values are shown as the difference between the fluorescence values for each phage and the fluorescence values for a negative control (dialysis buffer diluted in MEM), which were then normalized to the fluorescence of assayed supernatants from cells that received lysis buffer. Staurosporine (1 µM) was used as an apoptosis positive control. Error bars indicate SEM for three independent biological replicates.

To validate that phages do not cause loss of epithelial integrity, we measured transepithelial electrical resistance (TEER) during 48 h of phage treatment (Fig. 4B). Cell cultures incubated with phages maintained TEER over the time course of treatment. To verify that phages do not cause cell death, we evaluated phage cytotoxicity by measuring the release of the intracellular enzyme glucose-6-phosphate dehydrogenase (G6PD) into the apical media (Fig. 4C). We found that phage treatment was not cytotoxic in our CF AEC model, as it did not induce G6PD release. In summary, phages do not kill mammalian cells but do attach to areas where epithelial reorganization has occurred, which are often abundant in the airways of pwCF (32).

### Phages penetrate into the airway epithelium and their distribution is phage-specific

In our imaging experiments, we observed that phage localization was not solely distributed in the apical regions, and some phage particles were located throughout the epithelium. To complement these observations, we analyzed the depth distribution of phages in CF AECs after 1 and 24 h of phage treatment (Fig. 5A to Fig. 5D). At 1 h, we found that phage distribution across the epithelium was dissimilar for phages of different morphology (Fig. 5A and Fig. 5B). Phages OMKO1 and LPS-5 distributed all along the depth of the epithelium, with a mean distribution located around 50% of the total epithelial depth, and a similar proportion of particles above and below the mean depth. In contrast, phages PSA04 and PSA34 showed very different depth distributions after a 1 h incubation. Phage PSA04 was mostly distributed deep in the epithelium, although the mean depth was similar to phages OMKO1 and LPS-5. Phage PSA34 was mainly localized closer to the apical side, in agreement with the amount of aggregation observed on cells.

**Figure 5.**
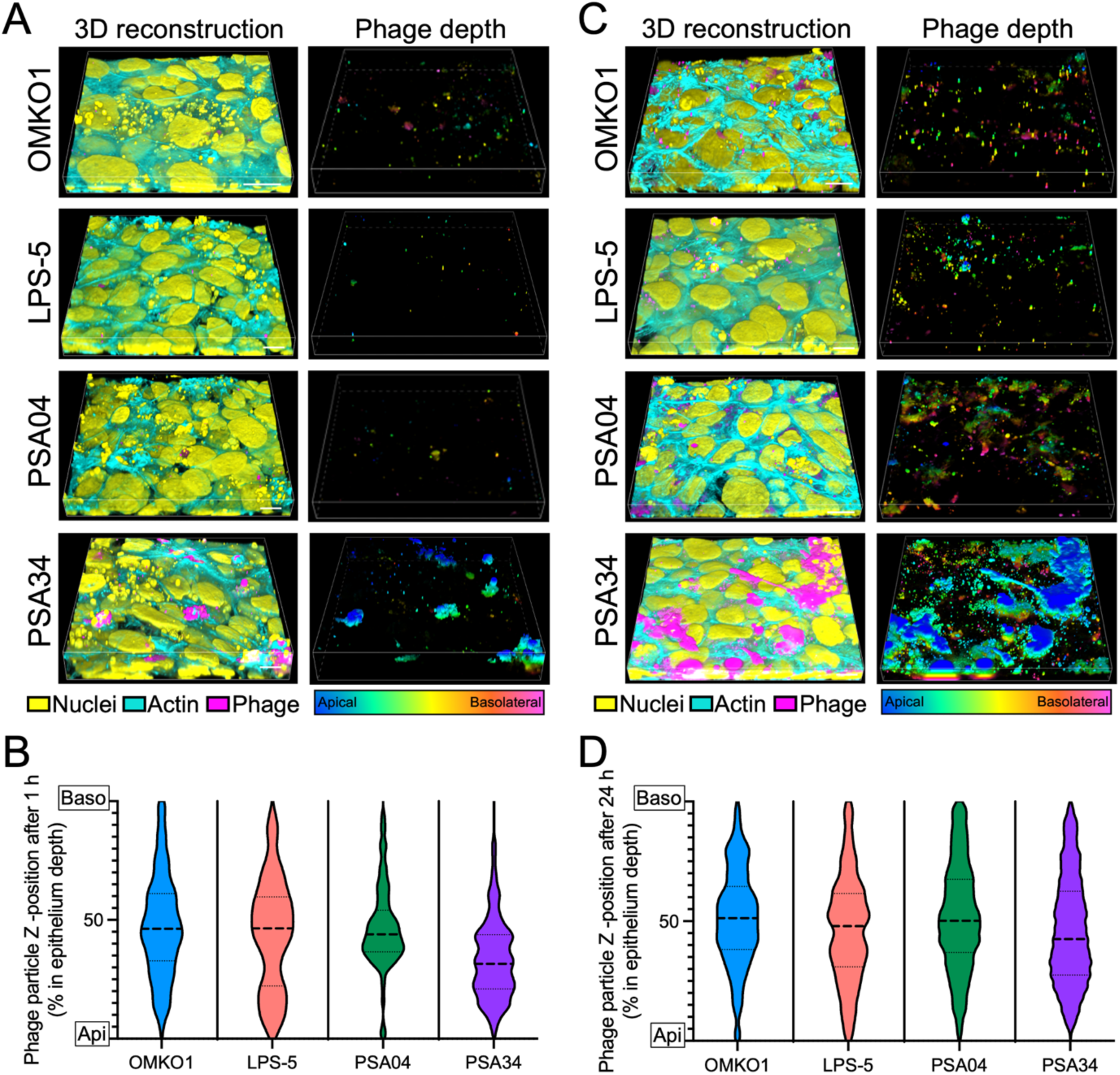
Phages penetrate into the airway epithelium and their depth is phage dependent. CFBE41o-cells were apically treated with fluorescently labeled phages OMKO1, LPS-5, PSA04, and PSA34 (magenta) at a concentration of 1 x 10^9^ PFU/ml in MEM pH-adjusted to 6.5 for 1 h (A and B) and 24 h (C and D). Cells were washed and fixed for confocal microscopy. Cells were stained with fluorescently labeled phalloidin (cyan) and Hoechst (yellow) to label actin and nuclei, respectively. (A and C) 3D reconstructions of confocal microscopy Z-stacks. Scale bar, 10 µm. TRITC-positive particles are shown as a depth-projected particles in a rainbow scale, with apically located particles in blue and basolateral-located particles in pink. (B and D) TRITC-positive particles were identified in 3D reconstructions using the 3D Object Measurement function in Nikon Elements. Their Z-location was plotted as percentage to the total depth of each Z-stack, and the thickness of the stacks was similar between experimental replicates. Plots show the combined depth from particles tracked from three independent confocal imaging stacks.

After 24 h of incubation, we discovered an increase in phage penetration, indicating that phage entry into the epithelium was time-dependent (Fig. 5C and Fig. 5D). Overall, phage penetration across the airway epithelium became deeper and more homogeneous between phages after 24 h, suggesting that phage morphology influences early interaction between phages and mammalian cells. Like the observations at 1 h, the location for phage PSA34 was skewed towards the apical side, whereas all other phages reached deeper locations. In summary, phage penetration into the airway epithelium is time-dependent and phage-specific. Myoviruses OMKO1 and LPS-5 showed a similar penetration pattern. Phage PSA04 penetrated deeper into the epithelium than the rest of the phages, whereas PSA34, the smallest of the tested phages, showed a tendency to localize apically.

To gain more insight into intracellular interactions between phages and mammalian cells, we conducted plaque assays to measure intracellular intact phages over time (10). As plaque assays only measure phages that can infect and produce progeny, this assay allows the quantification of phage particles that still have the capacity to replicate. Phages OMKO1, LPS-5, PSA04, and PSA34 were added apically to CF AECs. Intracellular intact phage was quantified at different times post-treatment. After 1 h, we found that phage internalization and phage survival in CF respiratory cells was generally poor, with less than 0.001% of the phage inoculum internalized (Fig. 6A). Phages LPS-5 and PSA04 trended towards higher titers than OMKO1 and PSA34, suggesting differences in phage internalization kinetics. At later time points, we found that phage LPS-5, the phage that showed the highest amount of internalized phage after 1 h of incubation, displayed intracellular titers that were 3-fold higher than the ones at 1 h, suggesting that internalization for LPS-5 reached saturation before the other phages on the panel (Fig. 6B). For phage OMKO1 and PSA04, titers at 24 h increased approximately 10-fold from the titers at 1 h. Phage PSA34 displayed the largest fold change at 24 h, with titers in CF AECs being approximately 25-fold higher than the titers at 1 h. In summary, phage morphology correlates with the amount of internalized phage over time, with the podovirus phage PSA34, the smallest of the tested phages, being the one whose internalization by the airway epithelium is favored after 24 h.

**Figure 6.**
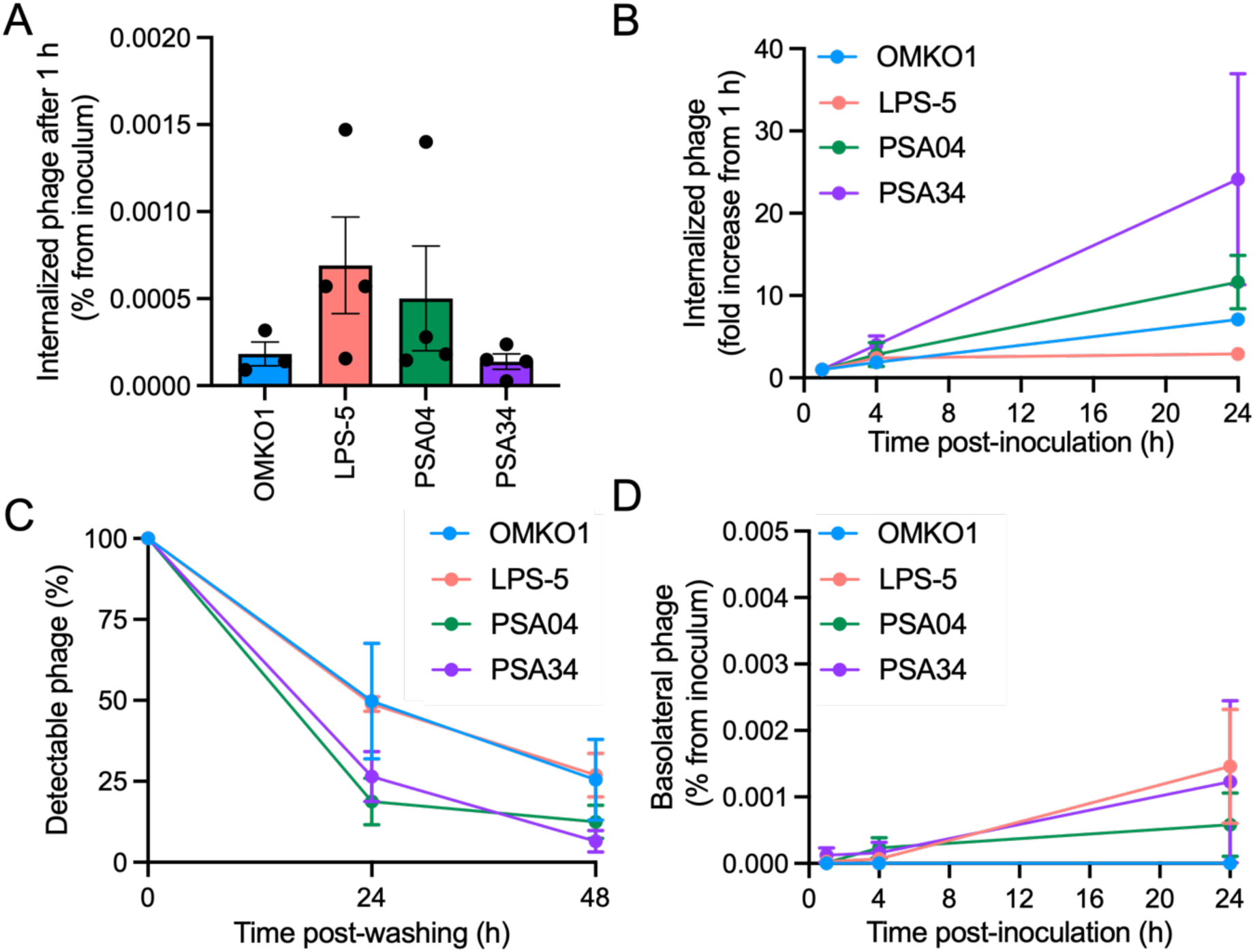
Phage internalization, degradation, and translocation into the basolateral space are phage specific. CFBE41o-cells were apically treated with phages OMKO1, LPS-5, PSA04, and PSA34 at a concentration of 1 x 10^9^ PFU/ml in MEM pH-adjusted to 6.5. (A) Internalized phages were quantified at 1 h post-treatment by plaque assay by measuring total associated phage in lysed cells versus extracellularly associated phage in intact cells. Results are depicted as the difference between lysed and intact cells. (B) Internalized phages were quantified at 1, 4, and 24 h post-treatment by plaque assay. Results at each time point were normalized to the titers at 1 h post-treatment. (C) After 24 h of phage treatment, cells were extensively washed to remove extracellular phages. At 0, 24 and 48 h following the extensive wash, internalized phages were quantified as described above. Results are plotted as percentage normalized to the amount of internalized phages at 0 h after the extensive wash. (D) Phage translocation into the basolateral space was measured by plaque assay at 1, 4, and 24 h post-apical phage treatment. Results are shown as the mean phage titers for at least three independent biological replicates. Error bars indicate SEM.

Next, we aimed to identify whether phages were degraded at different rates in mammalian cells. We incubated CF AECs with phages OMKO1, LPS-5, PSA04, and PSA34 for 24 h, followed by extensive washing to remove extracellular phages. We harvested cells at that time (time 0 h), 24 and 48 h later, and conducted plaque assays to quantify intact intracellular phages using a similar assay as the one described above (Fig. 6C). We found that phages OMKO1 and LPS-5 were degraded at similar rates, which were slower than the rates of degradation for phages PSA04 and PSA34. After 24 h, we detected phage titers equivalent to approximately 50% of the amount at the time of washing for phages OMKO1 and LPS-5, while we only detected 25% of active phage particles for phages PSA04 and PSA34 at the same time point. By 48 h, we detected 25% of phage titers for OMKO1 and LPS-5, versus approximately 10% for phages PSA04 and PSA34. These data suggest that within human respiratory epithelial cells, myoviruses are more stable than other types of phages. Phage degradation within cells is influenced by the intrinsic stability of each phage, as degradation assays of phages in fresh cell culture medium at 37°C in the absence of mammalian cells confirmed that myoviruses OMKO1 and LPS-5 remained viable for longer than phages PSA04 and PSA34 (Fig. S6).

Phages crossing to the basolateral space is a concern during phage therapy, as it increases the likelihood of phage entering the blood compartment and reaching other organs. To determine the epithelial barrier-crossing potential of each phage, we measured basolateral phage abundance after apical treatment in our CF AEC model (Fig. 6D). We found that phages differed in their ability to enter the basolateral compartment. Phage OMKO1, the largest phage in our panel, was unable to cross the epithelial layer and we failed to quantify infectious virus in the basolateral medium. However, we did detect an increase in basolateral phage over time for phages LPS-5, PSA04, and PSA34. For phage LPS-5, we detected higher titers in the basolateral space after a 24 h treatment compared to the intracellular titers, even as cells maintained TEER. This result suggests that the airway epithelium facilitates the translocation of LPS-5 from the apical to basolateral compartment. We detected approximately 50% of PSA34 titers in the basolateral space compared to intracellular titers, suggesting that PSA34 can cross the basolateral layer but it might not be as efficient as phage LPS-5. For phage PSA04, basolateral phage titers were approximately 15% of the intracellular titers, suggesting that phage translocation for PSA04 is poor. Overall, our results indicate that factors such as phage morphology, size, and aggregation affect phage-mammalian interactions, and thus, phage dynamics with the human host are likely specific for each type of phage.

### Airway epithelial cells respond to phages by increasing expression of cytokine signaling genes

We next sought to determine whether respiratory epithelial cells can sense and respond to purified lytic phages. We conducted RNA sequencing of RNA derived from phage-treated CF AECs. To enrich for phage-positive cells, CFBE41o-cells were exposed to fluorescently labeled OMKO1, LPS-5, PSA04, and PSA34 for 24 h and were then subjected to cell sorting using flow cytometry. We found that all tested phages induced transcriptional changes, with a core set of 12 genes significantly altered in all conditions assessed (Fig. 7A). All 12 genes encode proteins involved in cytokine signaling in the immune system, indicating that the phages were sensed as pathogen-associated molecular patterns (PAMPs) by the respiratory epithelium. The myoviruses OMKO1 and LPS-5, phages that showed better stability and the least aggregation on the airway epithelium, induced fewer changes in gene expression than the siphovirus PSA04 and the podovirus PSA34. Overall, phage treatment of CF AECs led to changes in expression of less than 2% of the evaluated genes, indicating that respiratory cells tolerate phage treatment without extensive transcriptome alterations. Pathway analysis revealed that the pathway with the highest combined Z-score was the pathway involved in pathogen-induced cytokine storm signaling, supporting our previous observation that the core set of differentially expressed genes for all phage-treated cells involved cytokine signaling in the immune system (Fig. 7B). Other upregulated pathways with high combined Z-scores were also related to the immune system, such as the multiple sclerosis signaling pathway and the macrophage classical activation signaling pathway, as well as cytoskeleton rearrangement pathways, including the FAK signaling pathway.

**Figure 7.**
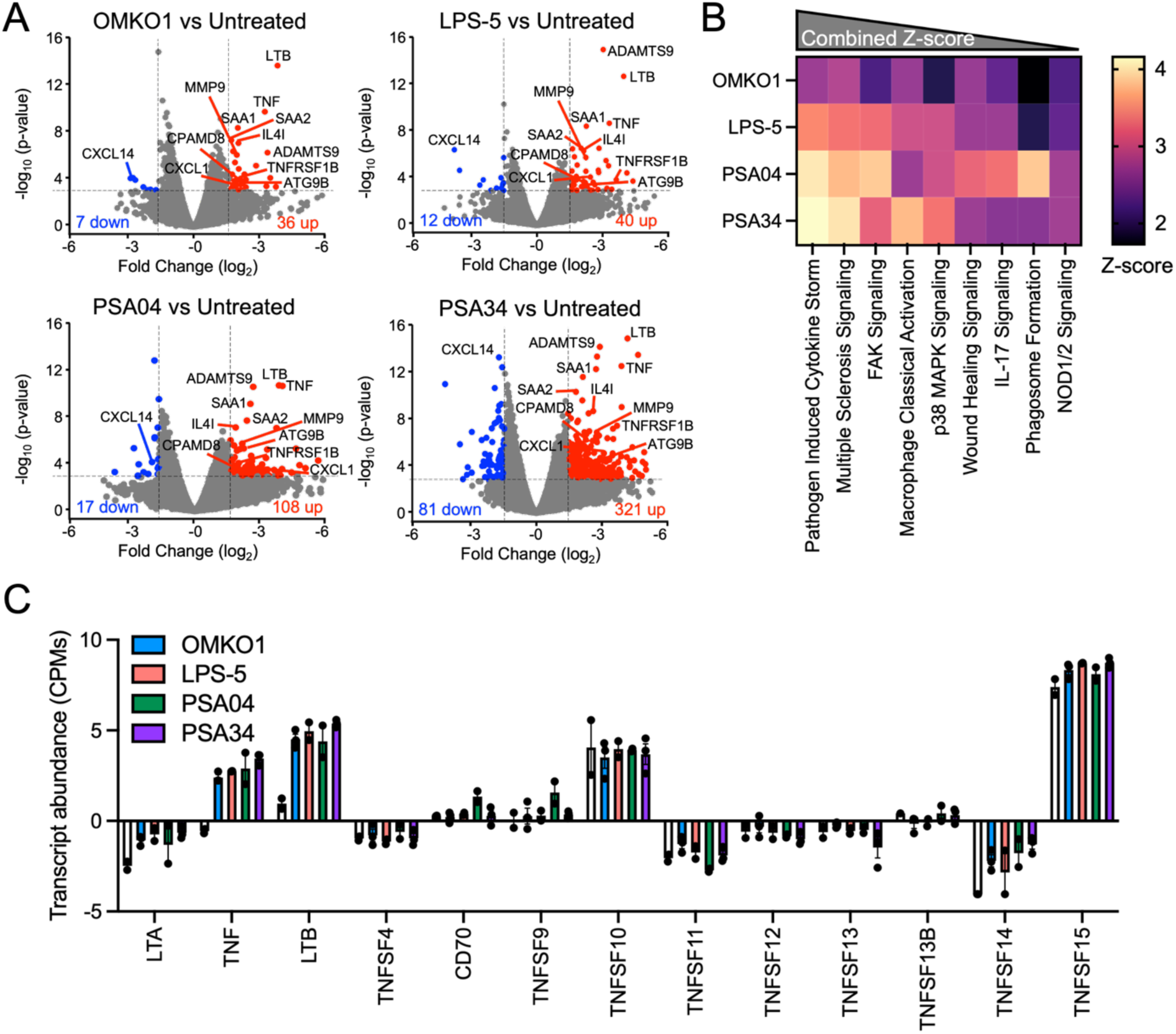
Airway epithelial cells change their transcriptome in response to therapeutic phages. CFBE41o-cells were apically treated with fluorescently labeled phages OMKO1, LPS-5, PSA04, and PSA34 at a concentration of 1 x 10^9^ PFU/ml in MEM pH-adjusted to 6.5 for 24 h. Cells were enriched for phage-positive cells using flow cytometry and RNA was sequenced. Following alignment to the human genome and normalization, differentially expressed genes were determined using EdgeR. (A) Volcano plots highlighting upregulated (red) and downregulated (blue) genes between phage-treated and untreated cells, using a fold change cutoff of +/- 1.5 and p<0.001. Depicted are genes that are differentially expressed in all phage-treatment conditions. (B) Ingenuity pathway analysis (IPA) for each set of differentially expressed genes. Heatmap depicts activation Z-scores, which predict that the indicated pathway and upstream regulators are more active in phage-treated cells than untreated ones. (C) Normalized transcript abundance for genes belonging to the TNF-superfamily. Genes with a very low transcript abundance (below 10) were not included.

We then analyzed the expression of individual genes belonging to the pathogen-induced cytokine storm signaling pathway and found that the highest changes in gene expression for all phage-treated cells were in genes of the tumor necrosis factor (TNF) superfamily (Fig. S7). The TNF superfamily is a protein superfamily of transmembrane proteins that are soluble upon extracellular proteolytic cleavage, which results in them becoming cytokines. We discovered that transcript abundance for LTA, LTB, TNF, and TNFSF14 were upregulated in phage-treated CFB41o-cells compared to untreated cells (Fig. 7C). Taken together, these data suggest that phages are detected by the airway epithelium as pathogens, inducing transcriptomic changes in cytokine signaling pathways.

### Cytokine secretion by the airways is phage-specific

To determine whether the changes in transcription of immune-related genes resulted in changes in cytokine secretion, we treated CF AECs with our panel of phages and collected apical and basolateral secretions at 24 h post-treatment. Cytokines were measured by a Luminex bead-based assay (Fig. 8A). We found that a specific group of cytokines were increased upon phage treatment. Most changes occurred for apically secreted cytokines, suggesting that the airway epithelium defense program is highly polarized. Cells treated with phages PSA04 and PSA34 induced the highest amount of proinflammatory cytokine secretion, reflecting the wider transcriptomic changes elicited by these phages. Cytokines TNFSF13B, TNFSF8, and IL-8 were significantly increased in the apical supernatant of cells treated with PSA04 and PSA34, whereas IFN-λ1 was significantly increased in cells exposed to OMKO1 and LPS-5 (Fig. 8B). All phages in our panel induced basolateral secretion of IFN-β, another antiviral cytokine. These results suggest that the respiratory epithelium responds to lytic phages by secreting antiviral and pro-inflammatory cytokines.

**Figure 8.**
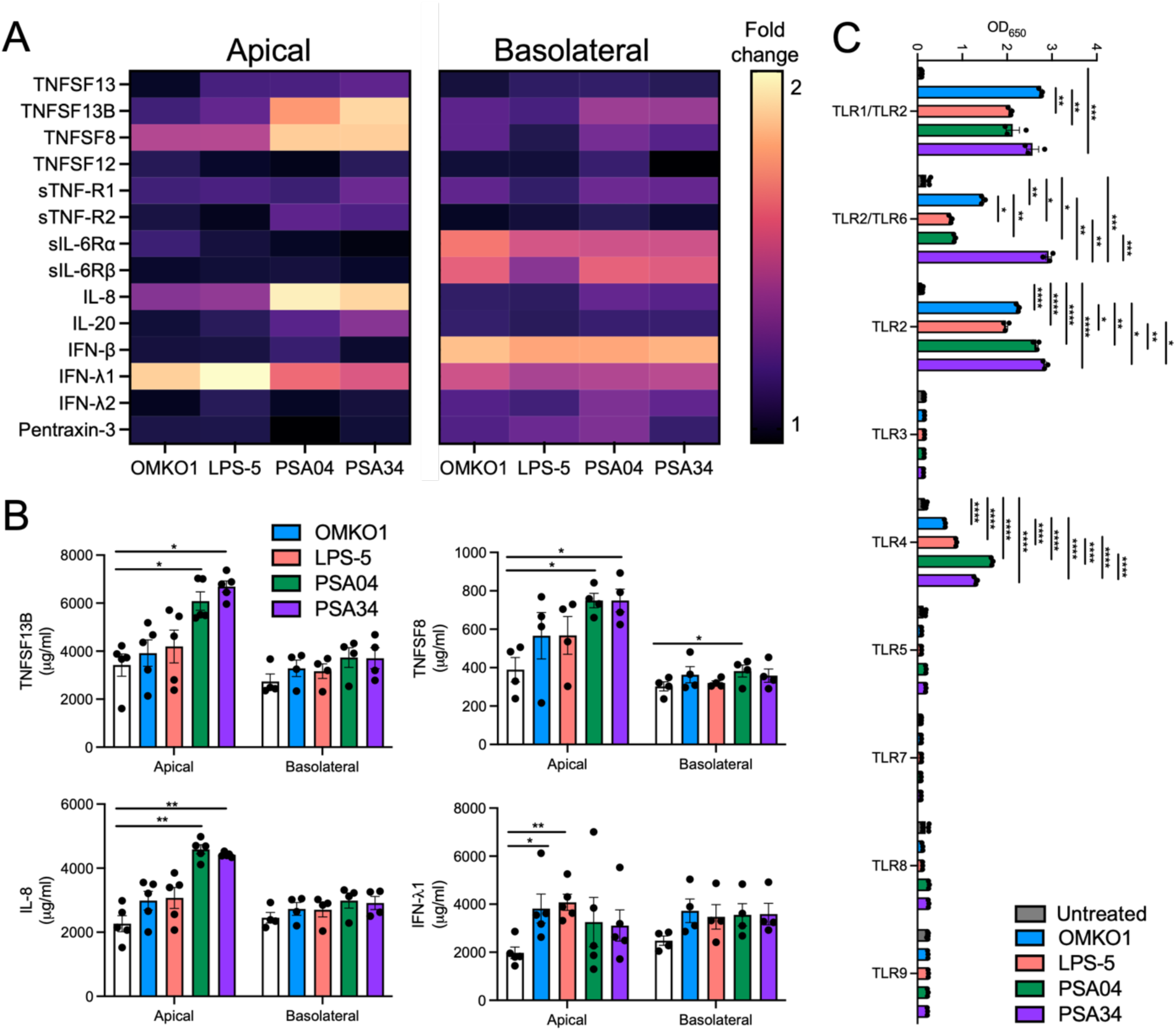
Airway epithelial cells sense and respond to phages by secreting pro-inflammatory cytokines. (A and B) CFBE41o-cells were apically treated with phages OMKO1, LPS-5, PSA04, and PSA34 at a concentration of 1 x 10^9^ PFU/ml in MEM pH-adjusted to 6.5 for 24 h. Supernatants and basolateral media were collected. Pro-inflammatory cytokines were measured using a multiplex cytokine array. Only cytokines above the limit of detection for all biological replicates are shown. (A) Heatmap depicting fold change between each phage-treated and untreated conditions. (B) Raw values for cytokines TNFSF13B, TNFSF8, IL-8, and IFN-λ1, which showed the largest changes between phage-treated and untreated conditions. (C) Toll-like receptor reporter HEK293 cells were treated with OMKO1, LPS-5, PSA04, and PSA34 at a concentration of 1 x 10^10^ PFU/ml for 24 h, followed by reporter activity measurements. Results are shown as raw absorbance values at 650 nm. Error bars indicate SEM for at least three independent biological replicates. *p<0.05, **p<0.01, ***p<0.001, ****p<0.000.1.

Because of the changes in transcription of immune related genes and the increase in cytokine secretion in response to phages, we next evaluated whether our panel of phages could engage pattern recognition receptors using a reporter epithelial cell line. HEK293 cells, which endogenously express little to no Toll-like receptors (TLRs, (33)), were transfected with different plasmids expressing human TLRs and a reporter gene to detect TLR activation. Phages were added to these reporter cell lines, and reporter activity was measured 24 h later (Fig. 8C). We found that all tested phages activated TLRs 1, 2, 4, and 6, suggesting that the immune responses detected at a transcriptomic and protein level were the result of PAMP recognition. In summary, phages with therapeutic potential engage extracellular TLRs and induce an increase in specific antiviral and proinflammatory cytokines. These responses are specific for each phage, and thus, should be taken into consideration when choosing phages for therapeutic treatment.

## DISCUSSION

The demand for alternative therapeutic treatments for bacterial infections is rapidly rising. Phage therapy offers several advantages compared to traditional antimicrobials, such as specificity and self-amplification. As a result, bacteriophages are increasingly being used in therapy to treat MDR infections. Even though they are reported as safe and the number of adverse events is often low, there are several gaps in knowledge regarding their interactions with the human host that are important to address as the number of phage therapy clinical trials grows. While phages do not replicate inside mammalian cells, high concentrations of phages come in close contact with mucosal surfaces during treatment. Here, we provide evidence that bacteriophage interactions with the airway epithelium are specific for phage type. These interactions are dependent on intrinsic phage properties, such as their morphology and aggregation, as well as extrinsic properties, such as characteristics of the physiochemical environment in the human host. Our data show that airway epithelial cells sense phages and mount transcriptomic changes that lead to the secretion of inflammatory cytokines, and that these responses are phage specific. Furthermore, our findings highlight the importance of expanding our knowledge regarding phage therapy, especially for the consequences in tissues of the human host. Performing assays that include mammalian cells during initial phage characterization will aid in the validation of this therapeutic alternative before it becomes more widespread. Our results demonstrate that there is heterogeneity in cellular responses to a group of different lytic *P. aeruginosa* phages with therapeutic potential, and thus, the effects of phages in the mammalian host cannot be generalized.

The ASL is the region of the airway mucosa where biofilms accumulate in many respiratory infections and, as such, is the location of interactions between phages and the mammalian epithelium during nebulized phage therapy. The pH of the ASL is critical to regulate innate immune defenses in the airways. Many chronic respiratory disorders are characterized by the dysregulation of the pH of the ASL, such as CF, COPD, and rhinosinusitis (29). Our study reveals that acidic pH influences phage interactions with airway epithelial cells, from increasing the number of phage particles associated with cells to favoring phage aggregation on the epithelium. This observation implies that in diseases characterized by an acidic mucosal environment, which occurs during inflammatory responses, phage-epithelial interactions are likely enhanced. Several protein receptors bind their ligands only at acidic pH, raising the potential of these receptors serving as attachment factors for phages (34, 35). Phages can bind glycans on the cell surface, in a similar way as mammalian viruses do (36), which is also affected by pH (37). The specific receptors that phages bind on mammalian cell surfaces remain to be elucidated. Understanding how epithelial-phage interactions are affected by the physiochemical environment of the airways is necessary, especially in CF, as its properties are altered after treatment with CFTR modulators (38–41). Inhaled corticosteroids and bronchodilators, which are used to treat CF and COPD, also reduce airway inflammation, and with that, affect airway pH, ionic composition, and mucin secretion. More studies are needed to fully understand the effects of the physiochemical environment on the tripartite relationship between phages, bacteria, and the human host in CF and other inflammatory disorders.

The use of phages with different morphologies allowed us to investigate the association between types of phages and their behavior in the airways. We discovered that phage internalization through the apical compartment is dependent on phage size and aggregation, especially in the first hour of incubation. The jumbo phage OMKO1 is internalized slowly, whereas the smaller phage LPS-5 enters cells more rapidly. PSA34, a phage that aggregates on cells, entered slowly during the first hour of treatment but it increased several fold by 24 h, suggesting that the entry of small phages into the airway epithelium is preferred over larger phages. PSA04, on the other hand, localized deeper in the epithelium compared to the other phages after the first hour of incubation. Overall, the amount of intact intracellular phage we detected was low compared to the starting inoculum, which diminishes some of the concerns regarding phage therapy having potential genotoxic effects in the human host (42, 43). Similar results have been obtained using other types of mammalian cells, such as gut and kidney epithelia (11, 12, 44). We found that virion stability is linked to phage morphology, with the myoviruses being more stable than the other tested phages, as it has been previously suggested (45). We observed that phage size is a factor that influences basolateral translocation; for example, the large OMKO1 phage was unable to translocate through the epithelium, whereas phages LPS-5, PSA04, and PSA34 could. These findings exemplify that different types of phages should be tested when examining interactions and activity in the human host, as their comportment with mammalian cells is phage specific.

Our work demonstrates that, despite the differences in internalization levels and degradation kinetics between different phages, all tested phages induced a core set of genes in airway epithelial cells. These genes encode for immune soluble mediators with functions in inflammation and wound healing. Interestingly, the responses that our panel of phages induces in CF AECs are distinct from what occurs after exposure of keratinocytes to Pf phage, a filamentous temperate phage produced in *P. aeruginosa* biofilms that is commonly found as a prophage in clinical isolates from pwCF (46–48). In keratinocytes, Pf phage promotes an epithelial response that prevents cellular migration and wound healing that might explain why skin infections with Pf phage-harboring *P. aeruginosa* are hard to heal (46). In the CF lung, tissue repair mechanisms are altered, with a reduction in cell migration and proliferation, and an increase in structural remodeling that leads to tissue thickening (32). Our data suggest that phage therapy could have effects beyond the capacity of phages to kill bacteria. In CF, an increase in wound healing could be beneficial, as it could restore some of the extracellular matrix components that are lost as the disease progresses (49). Transcriptomic analyses of CF cells treated with highly effective modulator therapy (HEMT) are needed to determine if the effects of phage therapy in wound healing pathways are dependent on CFTR function.

We found a range of epithelial immune responses induced by our panel of phages. Considering that our phage preparations are mostly devoid of bacterial components, the specific phage ligands that are responsible for immune stimulation remain to be determined. We found that our panel of phages engages TLRs that are often associated with extracellular pattern recognition (50). Although traditionally these receptors have been described as proteins that detect bacterial components, multiple lines of evidence indicate that TLRs are critical in damage-associated molecular pattern (DAMP) recognition and mediate sterile inflammation (51). TLRs also sense viral capsids through the detection of repeating subunit patterns (52) or specific structural proteins (53), and thus, we propose that similar mechanisms are involved in phage recognition by respiratory epithelial cells. Protein aggregation also triggers TLRs, as it has been demonstrated for endogenous proteins, such as some antibodies and amyloids (54). We observed that the respiratory epithelium released IFN-β in the basolateral space in response to all tested phages, indicating that an antiviral immune response was mounted. However, apical cytokine secretion was phage specific, with myoviruses OMKO1 and LPS-5 inducing an immune profile that was different from that induced by phages PSA04 and PSA34. OMKO1 and LPS-5 induced the secretion of IFN-λ1, another antiviral cytokine, whereas PSA04 and PSA34 stimulated the secretion of pro-inflammatory cytokines TNFSF13B, TNFSF8, and IL-8. Interestingly, these immune responses differ from those observed in mammalian cells treated with Pf phage, which induces an antiviral immune response, while preventing the secretion of TNF (55). Taken together, these collective observations reveal that cytokine responses to phages in the human host are phage specific, supporting the provision that phage-mammalian cell interactions should be considered to ensure phage therapy safety and optimal outcomes.

Our findings regarding immune responses to phages by the airway epithelium have important implications. Considering that different phages induce unique immune profiles, generalizations regarding phage effects in the human host cannot be made. This observation should be considered when choosing phages for therapeutic application, as immune polarization could either enhance or dampen phage therapy efficacy. For example, in a murine model of *S. aureus* infection, phage therapy improved wound healing in an independent manner to its direct effect on decreasing bacterial load (56). Such an effect could be detrimental in cases of fibrosis, where an increase in fibroblast activity and deposition of extracellular matrix impairs tissue function. In our case, phages such as OMKO1 and LPS-5, could be used as adjuvants to potentiate the effect of antiviral drugs (57, 58). Phages like PSA04 and PSA34, which increase IL-8 secretion, could be considered for most infections where neutrophil function is intact. However, they might not be optimal for pwCF, where increased neutrophil responses contribute to disease pathology (59). Phage therapy is already a personalized therapeutic treatment, as it requires the identification of a specific phage that lyses a patient’s unique bacterial isolate. Rational design for phage therapy has focused on the selection of phage cocktails that reduce the potential for phage resistance evolution (60). Our study suggests that immune response profiling should also be considered when choosing phages for a particular patient.

Our work has limitations that are important to address in future studies. Even though we included four phages with different physical properties that encompass the range of morphologies used in therapeutic phages, we cannot ensure that they represent the whole diversity of lytic phages. Developing high throughput assays using mammalian cells that assess phage-mammalian cell interactions and immune responses could improve the selection of phages that provide the most beneficial response in patients. The experiments conducted in this research were done using CFBE41o-cells cultured at air-liquid interface (ALI), which differentiate to a mucociliary phenotype, recapitulating the airways (61). However, other cell types might interact with phages in a different manner, as it is known that endocytic mechanisms are distinct in polarized cells (62). Additionally, our experiments did not address the effect of mucins in phage-epithelial interactions. Mucins are components of the ASL that are hypersecreted in CF. Phage binding to mucins has been suggested to be beneficial for human hosts, as it prevents phage diffusion and enhances potential interactions with their bacteria (63). The capacity of phages to bind to mucins is also specific to each phage and will likely impact the different types of phage associations (64). Our study focuses on mucosal innate immune responses, but understanding adaptive immune responses to phages is a critical step to ensure the efficacy of this therapeutic. Many treatment regimens involve consecutive doses over weeks or months and many studies have suggested that phage components are the target of mammalian antibodies (65–67). Finally, experiments involving phages, mammalian cells, and model microbiomes will be important to elucidate the effect of phages in their bacterial hosts, as well as bystander bacteria, in a more physiological context for the CF airways.

In the United States, the FDA recommends that phage source, endotoxin content, sterility, and lytic activity against the patient’s bacterial strains, to be provided when choosing a phage (68). However, these recommendations do not include assays involving mammalian cells or tissues. The results of our research, as well as the research of others (69), and the clinical cases that have reported adverse effects of phage therapy (8), lead us to suggest that these recommendations should be expanded to include documentation regarding phage-mammalian interactions, at least in non-emergency use situations. Assays like the ones conducted in our study could provide information that would help steer between choices of phages that have equivalent lytic activity. More research is needed to fully understand the phage determinants that trigger mammalian cell responses, as we and others have shown that phage preparations mostly devoid of LPS still can be recognized as pathogens.

## MATERIALS AND METHODS

### Cell culture

CFBE41o-cells (obtained from J. P. Clancy, Cincinnati Children’s Hospital, USA), a human CF ΔF508/ΔF508 airway epithelial cell, were cultured in Minimal Essential Media (MEM, Gibco, Thermo Fisher Scientific) supplemented with 10% of heat-inactivated fetal bovine serum (FBS, GeminiBio), 2 mM L-glutamine (Corning), and 5 U/ml penicillin-5 mg/ml streptomycin (Corning) in adherent cell culture conditions at 37°C with 5% CO_2_. To be used in experiments, CFBE41o-cells were seeded onto Transwell inserts (Costar) at a density of 1.5-2.5 x 10^5^ cells/insert for 6.5 mm inserts, 4-5 x 10^5^ cells/insert for 12 mm inserts, and 1.5-2 x 10^6^ cells/insert for 24 mm inserts. At 48 h, the apical media was removed to culture cells at an air-liquid interface (ALI). Basolateral media was replaced every other day until the time of the experiment. Cells were used between 2-3 weeks of culture, and after confirming transepithelial electrical resistance (TEER) to be over 300 Ω*cm^2^. All treatments to CF AECs were conducted apically in 100, 250, and 500 µl of MEM for 6.5-, 12-, and 24-mm inserts, respectively. The identity of CFBE41o-cells was confirmed by short tandem repeat profiling (University of Arizona Genetics Core, USA). Cells were routinely confirmed to be mycoplasma free by conducting PCRs using a mycoplasma detection kit (SouthernBiotech).

### Bacterial strains and culture conditions

*P. aeruginosa* laboratory and clinical strains (Table S2) were grown in lysogeny broth (LB, Fisher Scientific) at 37°C for 16 h with shaking at 250 rpm or on LB agar (Fisher Scientific).

### Preparation and propagation of bacteriophages

Phages PSA04 (GenBank: MZ089728.1) and PSA34 (GenBank: MZ089739.1) were received from the University of Pittsburgh (gift from Dr. Daria Van Tyne) and phages OMKO1 (GenBank: ON631220.1) and LPS-5 (FASTA file can be shared upon request) from Yale University (gift from Dr. Paul Turner) and plated for plaque purification using the double-layer agar method as described below. Plaques were picked using a sterile pipette tip 24 h later and used to infect their respective phage hosts in 5 ml cultures. Cultures were filtered 16 h later using 0.45 and 0.22 µm syringe filters with a polyethersulfone (PES) membrane (ThermoFisher Scientific), and the flow through was tittered by plaque assay as described below. Phage lysates were stored at 4°C and used for large scale phage purifications in 1 L cultures. *P. aeruginosa* strains PAO1 (for phages OMKO1 and LPS-5), DVT423 (for PSA04), and DVT411 (for PSA34) were grown at mid-log phase and infected with phages at an MOI of 0.005 PFU/cell for at 37°C for 16 h with shaking at 250 rpm in LB medium supplemented with 0.001 M of CaCl_2_ and MgCl_2_. Bacteria were lysed with 1% chloroform and removed by centrifugation at 5,500 x g for 20 minutes (min). Supernatants were treated with 1 µg/ml of DNase I (Sigma) and RNase A (ThermoFisher Scientific) at 37°C for 1 h, followed by vacuum filtration using a 0.45 µm filter and 0.22 µm filter with PES membranes (ThermoFisher Scientific). Supernatants were concentrated using Centricon Plus-70 centrifugal filter units with a membrane pore size of 100,000 Da (EMD Millipore) according to the manufacturer instructions. Centrifugations were repeated until supernatants were 100-fold more concentrated compared to the initial volume.

Supernatants were loaded onto cesium chloride (CsCl) gradients (densities: 1.70, 1.50, 1.45, and 1.33 g/ml), that were prepared in SM buffer (70) and centrifuged at 90,000 x g at 4°C for 16 h. Bands were collected using a 20-gauge syringe (Fig. S1), reloaded onto a similar CsCl gradient, and centrifuged again as before to increase phage purity. CsCl was removed by extensive dialysis using dialysis buffer (50 mM NaCl, 15 mM MgCl_2_, 10 mM Tris pH 7.4) and dialysis Float A Lyzer devices with a pore size of 10,000 Da (Sigma). Phage identity was confirmed by PCR using phage-specific primers (Table S1). Phage particle concentration was obtained by measuring absorbance at 269 and 320 nm and calculated using the (A_269_ -A_320_) x 6 x 10^16^ / number of bases per virion, as described (71). Preparations were tested for endotoxins using the Limulus amoebocyte lysate test (ThermoFisher Scientific) and bacterial DNA by amplification with 16S rRNA gene primers (72). All phage preparations used in experiments had endotoxin levels below 0.2 EU/ml in 1 x 10^8^ particles/ml.

### Phage DNA extraction, whole genome sequencing of phages, and genome assembly

Phages at a concentration of 1 x 10^12^ particles/ml were incubated with 600 mAU/ml of proteinase K at 56°C for 90 minutes. DNA was isolated using the DNeasy Blood & Tissue kit (Qiagen) according to the manufacturer instructions. Phage DNA was sequenced by the Microbial Sequencing and Analysis Center (MIGS, Pittsburgh, PA). Libraries were prepared with an Illumina DNA Prep kit with IDT 10bp UDI indices. Samples were sequenced on an Illumina NextSeq 2000 to obtain 2x151bp reads. Demultiplexing, quality checking, and adapter trimming was performed with bcl-convert (v3.9.3) (73). Samples were trimmed with trimmomatic version 0.36 (74). Phage genomes were assembled using SPAdes version 3.12.0 with the optional flag “--careful” (74) and annotated with prokka version 1.14.6 (75).

### Construction of bacteriophage phylogenetic tree

The bacteriophage tree was constructed with the Virus Classification and Tree Building Online Resource (VICTOR) server using whole genome concatenated amino acid sequences (76). Pairwise comparisons of bacteriophage amino acid sequences were generated with the Genome-BLAST Distance Phylogeny (GBDP) method (77) with the d_6_ formula (yielding average support of 34%), using settings recommended for prokaryotic viruses (76). Branch support was inferred from 100 pseudo-bootstrap replicates.

### PCR of phages

Phage DNA was extracted as described above, followed by quantification using a Nanodrop device (ThermoFisher Scientific). Phage DNA was amplified using phage specific primers (Table S1), which were designed using the Primer-BLAST tool (78), and GoTaq Green (Promega) according to manufacturer instructions. PCR products were visualized by electrophoresis on 0.5% Tris-acetate-EDTA (TAE) agarose gel at 100 V for 1 h using SYBR Safe DNA Gel Stain (Invitrogen).

### Fluorescent labeling of phages

Phages were diluted in 0.1 M of sodium bicarbonate at a concentration of 4 x 10^12^ particles/ml. Phage solutions were incubated with Alexa Fluor 555 NHS Ester (Succinimidyl Ester, ThermoFisher Scientific) at a concentration of 100 µM for 2 h at room temperature protected from light. Fluorescent phages were extensively dialyzed using dialysis buffer and dialysis tubing as described above. Phage fluorescence was confirmed by immunofluorescence microscopy before experimental use.

### Phage plaque assay

Quantification of infectious phages was conducted by the double-layer agar method. *P. aeruginosa* strains PAO1 (for phages OMKO1 and LPS-5), DVT423 (for PSA04), DVT411 (for PSA34), and clinical isolate strains from pwCF (Table S2), were grown at mid-log phase in LB medium as described above. Cultures were combined with fresh LB medium and “soft” agar (0.7%) at a ratio of 1:2:6, respectively. Solutions were poured over base LB agar (1.5%) and incubated overnight at 4°C. Phages were serially diluted in PBS supplemented with 0.9 mM CaCl_2_ and 0.5 mM MgCl_2_ and spotted on the double-agar plates. Plates with spotted phages OMKO1, PSA04, and PSA34 were incubated at 37°C and LPS-5 was incubated at room temperature for 16 h, and plaques were visualized and quantified on the next day.

### Transmission electron microscopy

Purified phages were prepared for imaging by transferring to a 200-mesh carbon coated grids, followed by negative staining with 1% uranyl acetate and blotting with filter paper. Grids were air dried and imaged using a JEOL 1400-PLUS 120KV transmission electron microscope at the Center for Biological Imaging at the University of Pittsburgh (Pittsburgh, PA), operated at 80kV. The International Committee on Taxonomy of Viruses (ICTV) guidelines were used for morphology classification (79). Length measurements for phage capsids and tails were obtained using ImageJ. At least three independent images were used, and the reported length corresponds to the mean length from those measurements.

### Phage killing assays in liquid culture

Mid-log phase cultures of *P. aeruginosa* strains PAO1, DVT423, and DVT411 grown in LB broth were OD_600_ normalized to 0.05 and seeded onto flat bottom, non-adherent, clear 96 well-plates. Phages OMKO1 (PAO1), LPS-5 (PAO1), PSA04 (DVT423), and PSA34 (DVT411) were added at an MOI of 0.01 PFU/bacterium and incubated at 37°C with shaking every 5 minutes. Bacterial growth over time was monitored by measuring absorbance at an optical density of 600 nm over 16 h using a SpectraMax M2 microplate reader (Molecular Devices).

### Biofilm quantification

Experiments that quantified abiotic bacterial biomass were done by growing *P. aeruginosa* strains PAO1 (for phages OMKO1 and LPS-5), DVT423 (for PSA04), and DVT411 (for PSA34) at mid-log phase in LB medium at 37°C for 18 h in 96-well plates. Bacteriophages were added at an MOI of 0.1 PFU/bacterium and incubated at 37°C for 6 h. After careful removal of supernatants, biomass was stained with crystal violet (41% crystal violet, 12% ethanol, 47% H_2_O) and quantified by solubilization in 30% acetic acid and measurement of absorbance at 550 nm on a SpectraMax M2 microplate reader (Molecular Devices).

Biotic bacterial biomass quantification was conducted by treating CFBE41o-cells with *P. aeruginosa* strains PAO1 (for phages OMKO1 and LPS-5), DVT423 (for PSA04), and DVT411 (for PSA34) at an MOI of 0.025 bacterium/cell in MEM pH 6.5 at 37°C for 1 h. Supernatants were removed and replaced with an equal volume of MEM pH 6.5 and incubated at 37°C for 16 h. Bacteriophages were added at an MOI of 1 PFU/bacterium and incubated at 37°C for 8 h. Supernatants were removed, and cells were washed with PBS supplemented with 0.9 mM CaCl_2_ and 0.5 mM MgCl_2_, followed by biomass solubilization using 0.1% Triton X-100 (Bio-Rad) in PBS. Biomass was centrifuged at 3,500 x g at room temperature for 3 minutes. Supernatants were removed and the pellet was resuspended in PBS supplemented with 0.9 mM CaCl_2_ and 0.5 mM MgCl_2_. Bacteria were serially diluted and plated on LB agar and incubated at 37°C for 16 h. Biomass was determined by counting CFUs.

### Phage-epithelial experiments using plaque assay

Phages associated with the airway epithelium were determined by treating CFBE41o-cells grown in Transwell inserts with phages OMKO1, LPS-5, PSA04, and PSA34 at a concentration of 1 x 10^9^ PFU/ml in MEM pH-adjusted to 6.5 or 7.5 at 37°C for 24 h. Cells were washed six times with PBS supplemented with 0.9 mM CaCl_2_ and 0.5 mM MgCl_2_, lysed with 0.1% Triton X-100 (Bio-Rad) in PBS, and phage infectious particles were quantified by plaque assay as described above. Phage internalization was determined by calculating the difference between total phage particles associated with the airway epithelium (lysed phage-treated cells) and extracellularly associated phage (intact phage-treated cells), as described (10). Briefly, to quantify phage internalization, phage treatments were conducted in duplicate wells. At the time of quantification, cells were detached enzymatically using TrypLE Select (ThermoFisher Scientific) from the Transwell inserts. Cells were centrifuged at 150 x g at 4°C for 5 min. Cells from the total phage well were resuspended in 0.1% Triton X-100 in PBS, whereas the other well was used to quantify extracellular phage by resuspending cells in PBS without detergent. Samples were serially diluted and used in plaque assays. Phage degradation assays were conducted by treating cells with phages as described above for 24 h. Cells were washed six times with PBS supplemented with 0.9 mM CaCl_2_ and 0.5 mM MgCl_2_ and returned to the 37°C incubator. Internalized phage was quantified at that time, 24, and 48 h later. Phage titers were calculated at each time and results were normalized to the titers obtained immediately after the initial wash step (0 h). Basolateral phage was quantified by collecting basolateral medium at 1, 4, and 24 h post-phage inoculation by plaque assay.

### Phage-epithelial experiments using immunofluorescence microscopy

CFBE41o-cells grown in Transwell inserts were treated with Alexa Fluor 555 fluorescent-OMKO1, LPS-5, PSA04, and PSA34 at a concentration of 1 x 10^9^ PFU/ml in MEM pH-adjusted to 6.5 or 7.5 at 37°C for 1 or 24 h. Cells were washed three times with PBS supplemented with 0.9 mM CaCl_2_ and 0.5 mM MgCl_2_, fixed with 4% paraformaldehyde (Electron Microscopy Sciences) in PBS, permeabilized with 0.1% Triton X-100 (Bio-Rad) in PBS, and stained with Alexa Fluor 647 phalloidin (ThermoFisher Scientific) at a concentration of 200 nM and Hoescht 33342 (ThermoFisher Scientific) at a concentration of 20 µM in PBS. Membranes were cut out using a dissecting scalpel (Electron Microscopy Science) and mounted on glass slides of 1.5 mm thickness (Fisherbrand) and covered with #1 thickness glass coverslips (Fisherbrand) using the ProLong Gold Antifade mounting medium (ThermoFisher Scientific). Images were obtained using a Nikon C2 confocal system and a Plan Apo 60X oil/1.40NA objective. Images were processed using Nikon Elements version 5.10, using the denoise.ai function. Phage particles were analyzed using 3D volumetric reconstructions and the 3D Object Measurement function in the Nikon Elements Analysis software. Particles under 0.05 µm^3^ were discarded from the analysis with the intention to reduce background noise. Particle penetration into the depth of the airway epithelium was determined by calculating the distance in the Z-axis from the top of each Z-stack to each particle over 0.05 µm^3^, with Z-stacks of similar dimensions between experimental replicates.

### Epithelial viability assays

To evaluate epithelial integrity following phage treatment, CFBE41o-cells grown in Transwell inserts were treated with phages OMKO1, LPS-5, PSA04, and PSA34 at a concentration of 1 x 10^9^ PFU/ml in MEM pH-adjusted to 6.5 at 37°C. At 0, 24, and 48 h following treatment, MEM was added to the apical compartment, incubated at 37°C for 30 minutes, and transepithelial electrical resistance was measured using a EVOM2 chopstick electrode set (World Precision Instrument STX2). Results are shown as the difference obtained in Transwell inserts containing cells versus blank Transwell inserts. Cytotoxicity from phages was analyzed by treating CFBE41o-cells grown in Transwell inserts with phages OMKO1, LPS-5, PSA04, and PSA34 at a concentration of 1 x 10^9^ PFU/ml in MEM pH-adjusted to 6.5 at 37°C for 24 h. Apical supernatants were collected and combined with resazurin as defined in the CyQUANT Cytotoxicity Assay Kit (ThermoFisher Scientific). Resofurin, which is proportional to the amount of G6P in the medium, was measured after 30 min of incubation by fluorescence (excitation at 540 nm, emission at 590 nm) using a Molecular Devices SpectraMax M2 plate reader. Staurosporine at a concentration of 1 µm (Selleckchem) was included as an apoptosis positive control. Results were depicted as the difference between phage-treated and untreated samples and normalized to the cytotoxicity values of cells treated with Cell-Lysis Buffer (ThermoFisher Scientific). Apoptosis measurements were conducted by visualizing phosphatidylserine translocation by confocal microscopy. Briefly, CFBE41o-cells grown in Transwell inserts were treated with phages for 24 h as described above. Cells were washed with PBS and incubated with Annexin V-FITC (ThermoFisher Scientific) in Binding Buffer (10 mM HEPES pH 7.4, 140 mM NaCl, 2.5 mM CaCl_2_) as indicated by the manufacturer instructions and incubated at room temperature for 15 min. Cells were washed with Binding Buffer, fixed with 4% paraformaldehyde in PBS, and processed for confocal microscopy as described above.

### Nanoparticle tracking analysis (NTA)

Phages OMKO1, LPS-5, PSA04, and PSA34 at a concentration of 1 x 10^8^ PFU/ml were incubated in minimal essential media pH-adjusted to 6.5 or 7.5 at 37°C for 24 h. Phages were analyzed in a NanoSight NS300 (Malvern Panalytical Ltd) using a syringe pump speed of 30 AU and a sCMOS camera set up to take three captures of 30 seconds per sample. Other parameters were arranged according to the manufacturer instructions.

### Cell sorting and RNA sequencing

CFBE41o-cells grown in Transwell inserts were treated with Alexa Fluor 555 fluorescent-OMKO1, LPS-5, PSA04, and PSA34 at a concentration of 1 x 10^9^ PFU/ml in MEM pH-adjusted to 6.5 at 37°C for 24 h. Cells were washed three times with PBS supplemented with 0.9 mM CaCl_2_ and 0.5 mM MgCl_2_, and incubated with TrypLE Select (ThermoFisher Scientific) at 37°C for 40 min to detach cells. Cells were collected, centrifuged at 370 x g at 4°C for 5 minutes, and resuspended in PBS supplemented with 2% FBS. Cells were strained through a 40 µm mesh size strainer (ThermoFisher Scientific) and flow sorted into Alexa Fluor 555-positive and negative populations using a BS LSR II flow cytometer at the Flow Cytometry Core at the Department of Immunology at the University of Pittsburgh. Cells were sorted into DNA/RNA Shield (Zymo Research) and submitted to Zymo Research for RNA extraction and total RNA sequencing. RNA extraction was conducted using the Quick-RNA Miniprep kit (Zymo Research), followed by RNA clean-up using the RNA Clean & Concentrator-5 kit (Zymo Research). Library preparation was done using the Zymo-Seq RiboFree Total RNA Library Kit (Zymo Research) and libraries were sequenced on an Illumina NovaSeq instrument with a sequencing depth of 30 million read pairs.

### RNA sequencing bioinformatic analysis

Raw reads quality was first confirmed using FastQC v.0.11.9 (80). Reads were trimmed for adapters and low-quality sequences using Trim Galore! v0.6.6 (81) and aligned to the reference human genome GRCh38/hg38 using STAR v2.6.1d (82). Reads mapping to exons were assigned using featureCounts v2.0.1 (83). Quality control was tracked using MultiQC v1.9 (84). Differential gene expression analysis was performed using EdgeR v3.42.4 (85). Low expression genes were removed (counts less than 10), data was normalized by the trimmed mean of M-values, and contrasts were conducted between each phage’s dataset versus the dataset of untreated samples. Volcano plots were generated using the EnhancedVolcano v3.17 (86). Genes were considered to be significant if the differential gene expression log_2_ fold change was greater than |1.5| and p ≤ 0.001. Pathway analysis was conducted using Ingenuity Pathway Analysis (IPA, Qiagen) using genes with differential gene expression log_2_ fold change greater than |1.5|.

### Cytokine analysis

Phages associated with the airway epithelium were determined by treating CFBE41o-cells grown in Transwell inserts with phages OMKO1, LPS-5, PSA04, and PSA34 at a concentration of 1 x 10^9^ PFU/ml in MEM pH-adjusted to 6.5 at 37°C for 24 h. Apical and basolateral media was collected, centrifuged at 370 x g at 4°C for 5 min, and human proinflammatory cytokines were quantified using the Bio-Plex Pro™ Human Inflammation Panel 1 (BioRad). Fluorescence was read using a MAGPIX System (Luminex) at the UPMC Children’s Hospital of Pittsburgh (Pittsburgh, PA). Cytokines that showed values below the limit of detection were discarded from the analysis.

### Pattern Recognition Receptor Screening

Toll-like receptor screening was conducted by seeding HEK-Blue hTLR2-TLR1, HEK-Blue hTLR2-TLR6, HEK-Blue hTLR2, HEK-Blue hTLR3, HEK-Blue hTLR4, HEK-Blue hTLR5, HEK-Blue hTLR7, HEK-Blue hTLR8, and HEK-Blue hTLR9 (InvivoGen) into 96-well plates. Phages OMKO1, LPS-5, PSA04, and PSA34 were added at a concentration of 1 x 10^10^ PFU/ml in PBS supplemented with 0.9 mM CaCl_2_ and 0.5 mM MgCl_2_ and incubated at 37°C for 24 h with 5% CO_2_. At the end of the incubation, NF-κB activity was monitored by the production of secreted embryonic alkaline phosphatase (SEAP) through the hydrolysis of HEK-Blue (InvivoGen). Optical density was read at 650 nm using a SpectraMax340 PC plate reader (Molecular Devices).

### Statistical analysis

Statistical analysis for *in vitro* and cell culture experiments was conducted using GraphPad Prism v.10, except for the RNA sequencing analysis that it was analyzed as described above. Cell culture experiments were conducted using at least three independent biological replicates, which were defined as independent experiments performed on separate days using different passages of CFBE41o-cells. RNA sequencing experiments were conducted using at least two independent biological replicates. Data are presented as mean ± the standard error of the mean (SEM), unless mentioned in the figure legends. Two-tailed Student’s t test or one-way ANOVA followed by Dunnett’s post-hoc test were used to define statistical significance.

### Data availability

RNA sequencing raw counts tables were deposited in NCBI’s Gene Expression Omnibus (GEO) and are publicly available at the time of publication.

## ACKNOWLEDGMENTS

We thank members of the Bomberger lab for their suggestions and critical review of this manuscript. We thank Roxanna Barnaby for aiding with the NTA experiments. We thank the CF Bioinformatics & Biostatistics Core (P30) at the Geisel School of Medicine at Dartmouth for their help analyzing RNA sequencing data. We thank Michelle Manni, PhD for her help with the MAGPIX System. We thank the personnel from the following cores for their experimental help: Center for Biological Imaging and the Flow Cytometry Core at the Department of Immunology at the University of Pittsburgh.

This work was supported by a Cystic Fibrosis Foundation (CFF) Postdoctoral Fellowship ZAMORA20F0 and NIH grant T32AI060525 to P.F.Z.; CFF grants ARMBRU19F0 and ARMBRU22F5 and NIH grant T32HL129949 to C.R.A; and CFF grant BOMBER21P0 to J.M.B. Conflict of Interest statement: PET is cofounder of Felix Biotechnology, Inc., a company that seeks to develop phages for human therapy. Yale University has an institutional conflict of interest related to this project (PET & JLK). Yale may receive financial benefit related to the therapy used in this protocol.

---

**Figure S1.**
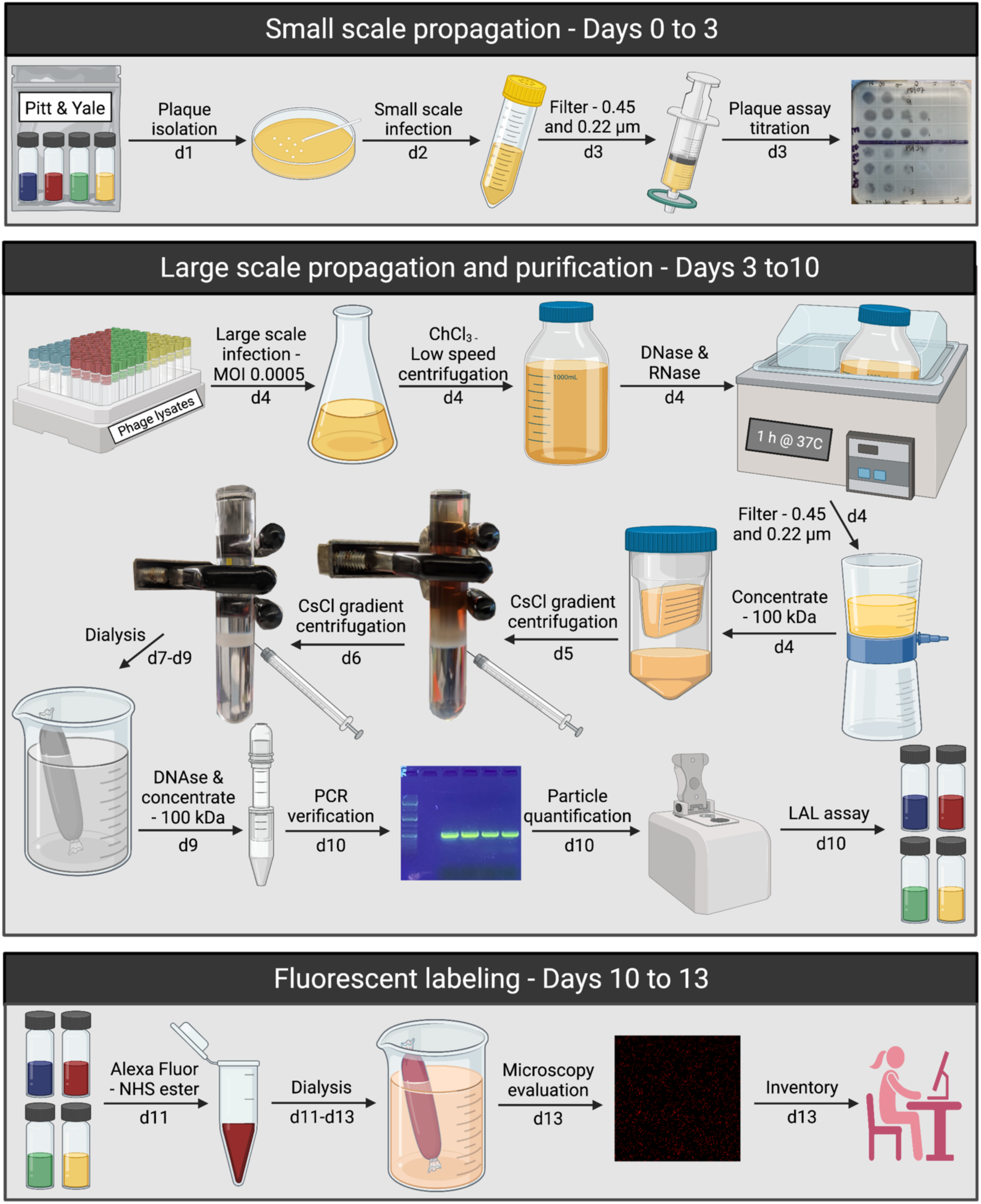
Protocol to prepare highly purified phages. Bacteriophages OMKO1, LPS-5, PSA04, and PSA34 were shipped at 4°C and used in a small-scale propagation to generate phage lysates that were stored at 4°C. Before large scale propagations, phages were re-tittered by plaque assay and used to infect large volume cultures of *P. aeruginosa* strains PAO1 (for phages OMKO1 and LPS-5), DVT423 (for PSA04), and DVT411 (for PSA34). Filtered and concentrated phage lysates were loaded onto CsCl gradients and phages were twice purified by gradient ultracentrifugation. Phage bands were collected, extensively dialyzed to remove CsCl, and phage identity was verified by PCR using phage-specific primers (Table S1). Phage particles were quantified by absorbance at 269 and 320 nm. Endotoxin levels were quantified using the limulus amoebocyte lysate (LAL) test. Phages to be used in microscopy experiments were fluorescently labeled with Alexa Fluor dyes and fluorescence was confirmed before experiments by microscopy. Figure was created with BioRender.com.

**Figure S2.**
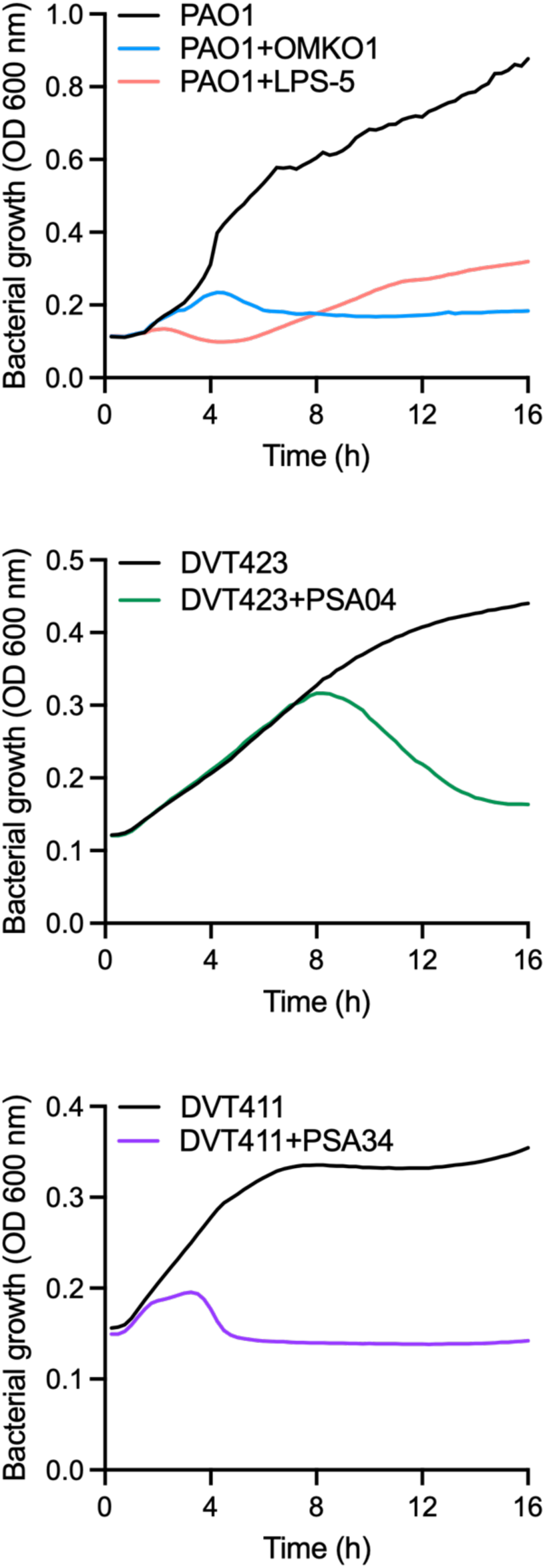
Phages show different killing kinetics in liquid culture. *P. aeruginosa* strains PAO1, DVT423, and DVT411 grown in LB broth were incubated with phages OMKO1 (PAO1), LPS-5 (PAO1), PSA04 (DVT423), and PSA34 (DVT411) at an MOI of 0.01 PFU/bacterium at 37°C. Bacterial growth over time was monitored by measuring absorbance at an optical density of 600 nm over 16 h. Results represent the mean from three independent experiments.

**Figure S3.**
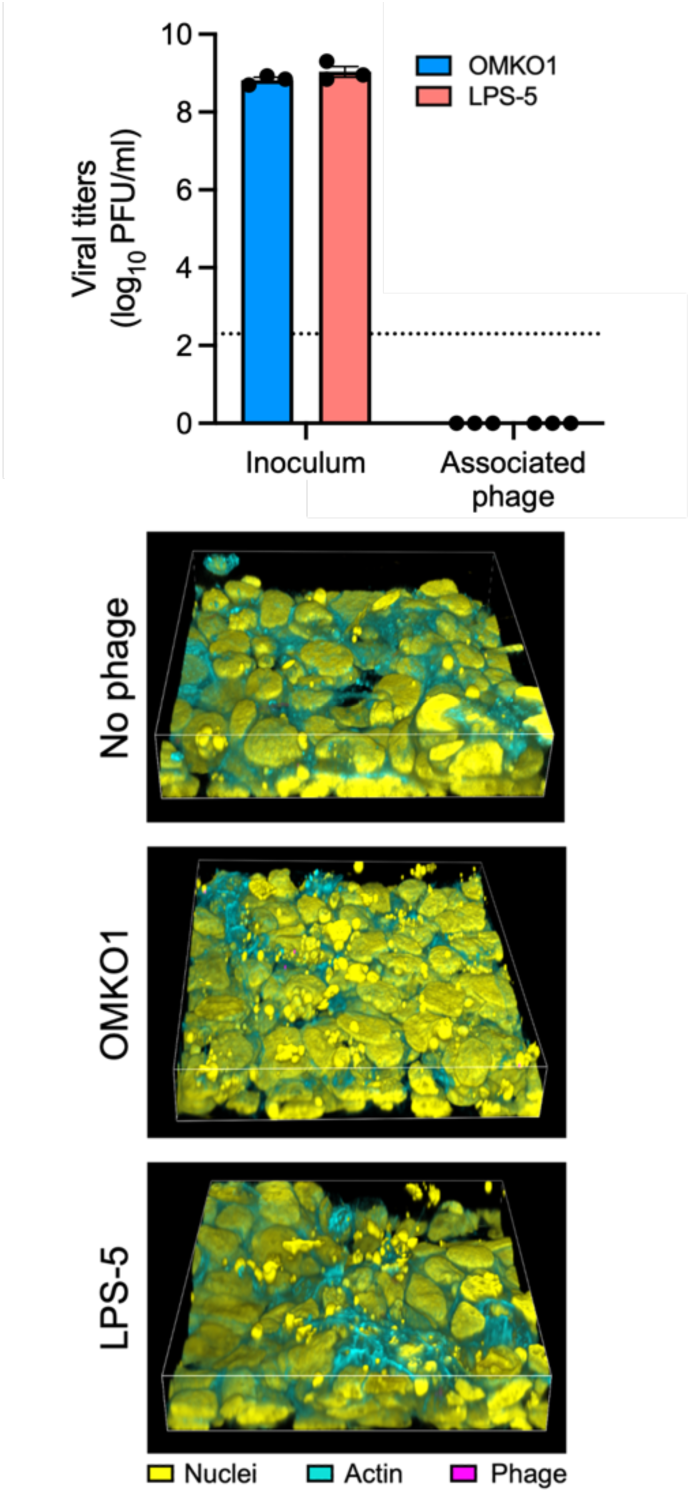
Phage background signal is minimal in the experimental association assays with epithelial cells. Unlabeled (top) and fluorescently labeled (bottom) OMKO1 and LPS-5 phages were incubated with CFBE41o-cells at an MOI of 1 x 10^9^ PFU/ml, following by immediate removal of the inoculum (0 h post-treatment). These phages were chosen for this experiment as they represent the phages with the highest and lowest internalized phage after a 1 h incubation (data from Figure 6). Inoculum and cell-associated phages were quantified by plaque assay (top). After inoculum removal, cells were fixed and processed for confocal microscopy (bottom). Images are shown as 3D reconstructions. LOD, limit of detection.

**Figure S4.**
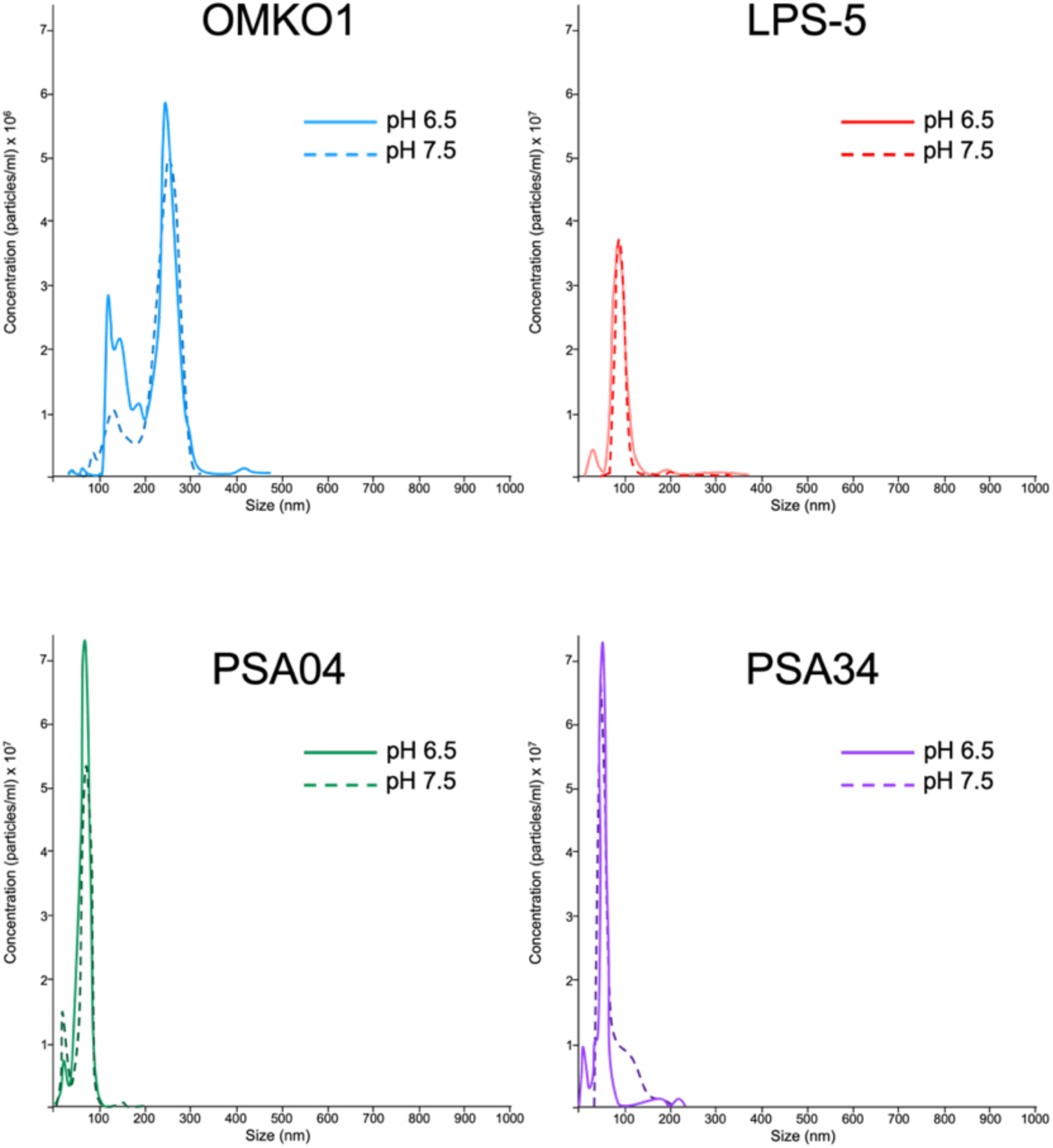
Acidic pH does not affect phage particle size in cell culture-free conditions. Phages OMKO1, LPS-5, PSA04, and PSA34 were incubated in minimum essential media at pH 6.5 or 7.5 at a concentration of 1 x 10^8^ PFU/ml at 37°C for 24 h. Particle dynamics was tracked using a nanoparticle analyzer

**Figure S5.**
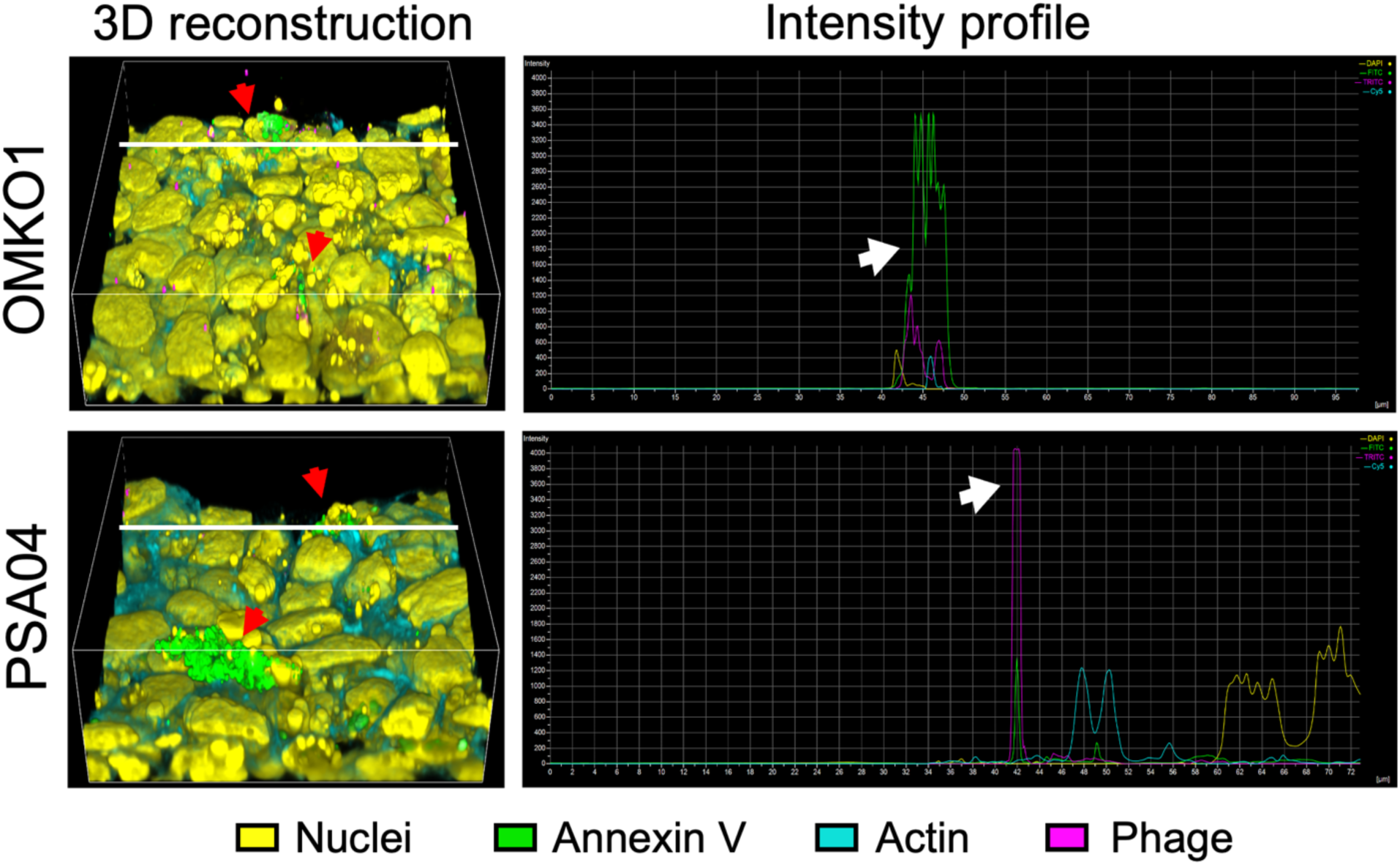
Phage particles locate in epithelial areas where apoptosis has ocurred. CFBE41o-cells grown in Transwell inserts were incubated with phages OMKO1 and PSA04 at an MOI of 1 x 10^9^ PFU/ml at 37°C for 1 h. These phages were chosen for this experiment as they represent phages with different interaction patterns with the airway epithelium. Cells were stained with annexin V to label apoptotic areas, fixed, and processed for confocal microscopy. Images show 3D reconstructions (left), red arrowheads indicate areas positive for annexin V staining. White line depicts the location in the XY dimension chosen for the intensity profiles (right), which portray fluorescence intensity over the X-axis. White arrows indicate areas of colocalization between annexin V and phage staining.

**Figure S6.**
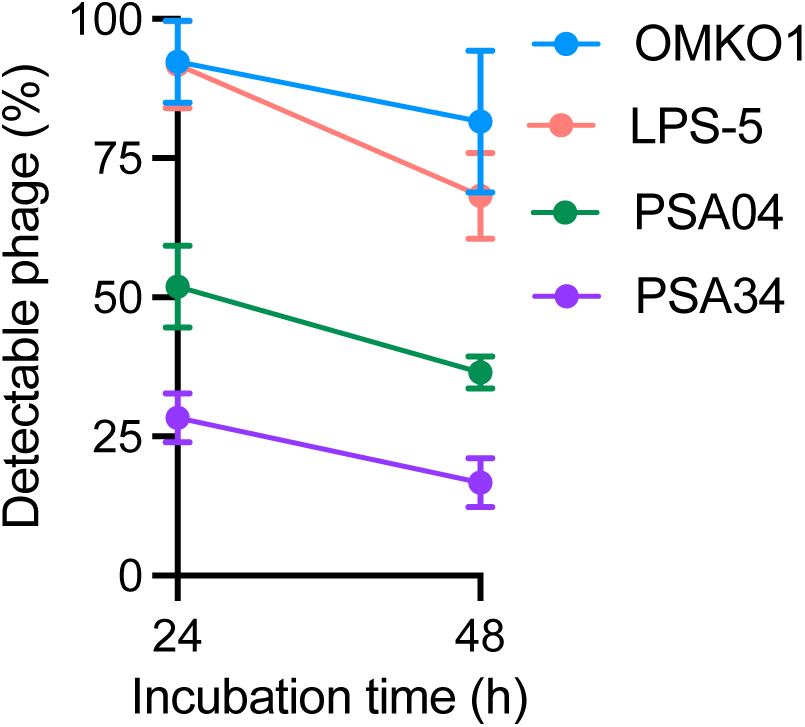
Phages have different physical stability in cell culture medium. Phages OMKO1, LPS-5, PSA04, and PSA34 were incubated in tissue culture media at a concentration of 1 x 10^9^ PFU/ml at 37°C for 48 h. Phages were tittered at 0, 24, and 48 h after the start of the incubation. Results are shown as percentage titers at each time point compared to the titers at 0 h. Results show the mean of three independent biological replicates, with error bars indicating SEM.

**Figure S7.**
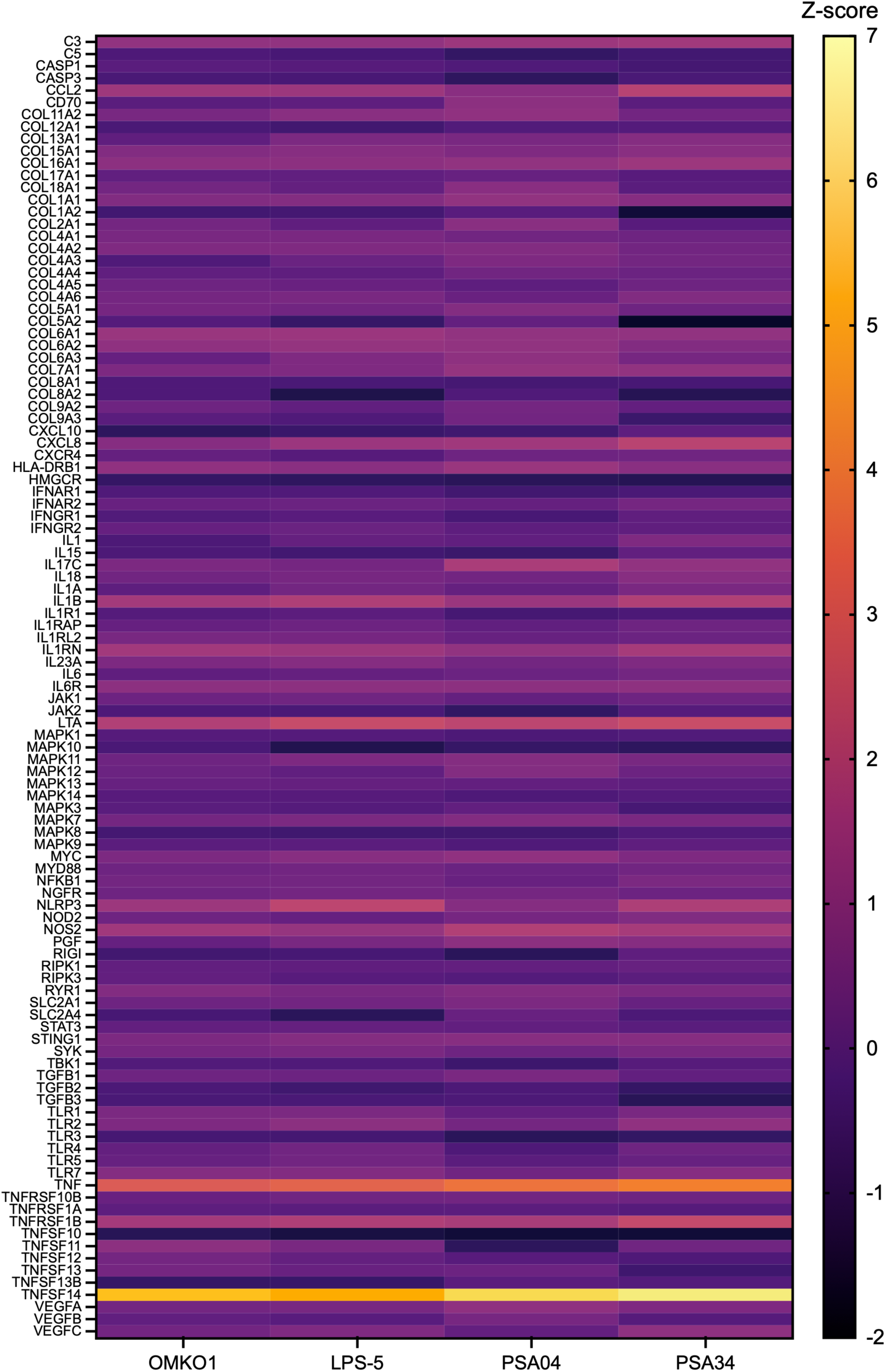
TNF superfamily genes are predicted to be activated following phage treatment. Individual Z-score values for genes in the “Pathogen Induced Cytokine Storm” pathway. Heatmap indicates Z-scores, as determined by the IPA z-score algorithm (87).

**Table S1.**
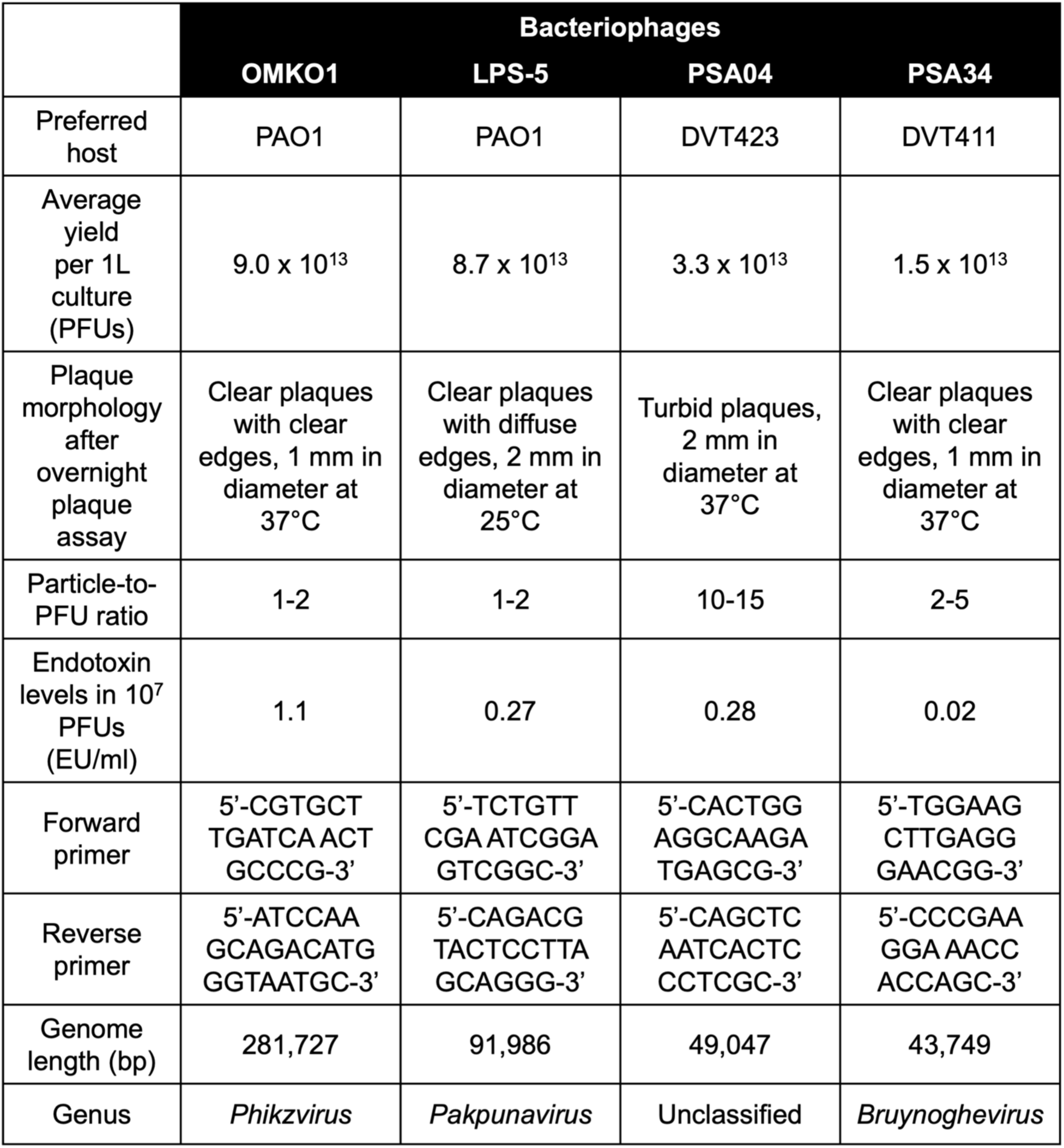
Summary of phage features.

**Table S2.**
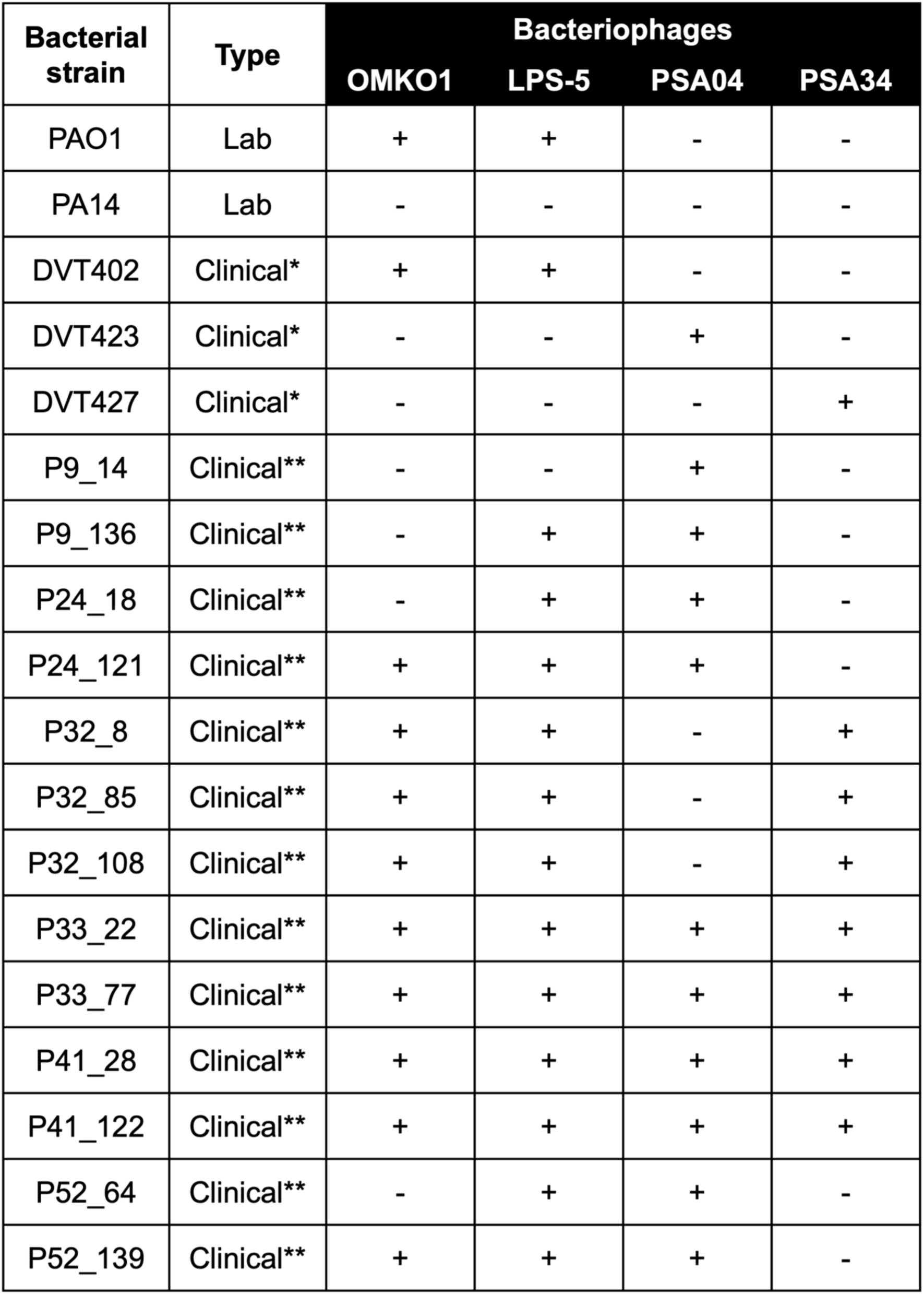
Phage killing activity over different *P. aeruginosa* strains. Laboratory and clinical isolate strains of *P. aeruginosa* were tested for susceptibility towards phages OMKO1, LPS-5, PSA04, and PSA34 by plaque assay. “+” indicates susceptibility, “-“ denotes resistance to the particular phage. *, clinical isolate strains obtained from CF sputum (24); ** clinical isolate strains obtained from CF sinuses (88).

## REFERENCES

1. Hatfull GF, Dedrick RM, Schooley RT. 2022. Phage Therapy for Antibiotic-Resistant Bacterial Infections. Annu Rev Med 73:197–211.

2. Hitchcock NM, Devequi Gomes Nunes D, Shiach J, Valeria Saraiva Hodel K, Dantas Viana Barbosa J, Alencar Pereira Rodrigues L, Coler BS, Botelho Pereira Soares M, Badaro R. 2023. Current Clinical Landscape and Global Potential of Bacteriophage Therapy. Viruses 15.

3. Petrovic Fabijan A, Iredell J, Danis-Wlodarczyk K, Kebriaei R, Abedon ST. 2023. Translating phage therapy into the clinic: Recent accomplishments but continuing challenges. PLoS Biol 21:e3002119.

4. Zaczek M, Gorski A, Weber-Dabrowska B, Letkiewicz S, Fortuna W, Rogoz P, Pasternak E, Miedzybrodzki R. 2022. A Thorough Synthesis of Phage Therapy Unit Activity in Poland-Its History, Milestones and International Recognition. Viruses 14.

5. Zaczek M, Weber-Dabrowska B, Miedzybrodzki R, Lusiak-Szelachowska M, Gorski A. 2020. Phage Therapy in Poland - a Centennial Journey to the First Ethically Approved Treatment Facility in Europe. Front Microbiol 11:1056.

6. Organization WH. 2022. Global antimicrobial resistance and use surveillance system (GLASS) report 2022. Geneva.

7. Fernandes P. 2015. The global challenge of new classes of antibacterial agents: an industry perspective. Curr Opin Pharmacol 24:7–11.

8. Liu D, Van Belleghem JD, de Vries CR, Burgener E, Chen Q, Manasherob R, Aronson JR, Amanatullah DF, Tamma PD, Suh GA. 2021. The Safety and Toxicity of Phage Therapy: A Review of Animal and Clinical Studies. Viruses 13.

9. Monteiro R, Pires DP, Costa AR, Azeredo J. 2019. Phage Therapy: Going Temperate? Trends Microbiol 27:368–378.

10. Lehti TA, Pajunen MI, Skog MS, Finne J. 2017. Internalization of a polysialic acid-binding Escherichia coli bacteriophage into eukaryotic neuroblastoma cells. Nat Commun 8:1915.

11. Bichet MC, Chin WH, Richards W, Lin YW, Avellaneda-Franco L, Hernandez CA, Oddo A, Chernyavskiy O, Hilsenstein V, Neild A, Li J, Voelcker NH, Patwa R, Barr JJ. 2021. Bacteriophage uptake by mammalian cell layers represents a potential sink that may impact phage therapy. iScience 24:102287.

12. Nguyen S, Baker K, Padman BS, Patwa R, Dunstan RA, Weston TA, Schlosser K, Bailey B, Lithgow T, Lazarou M, Luque A, Rohwer F, Blumberg RS, Barr JJ. 2017. Bacteriophage Transcytosis Provides a Mechanism To Cross Epithelial Cell Layers. mBio 8.

13. Kortright KE, Chan BK, Koff JL, Turner PE. 2019. Phage Therapy: A Renewed Approach to Combat Antibiotic-Resistant Bacteria. Cell Host Microbe 25:219–232.

14. Aslam S, Lampley E, Wooten D, Karris M, Benson C, Strathdee S, Schooley RT. 2020. Lessons Learned From the First 10 Consecutive Cases of Intravenous Bacteriophage Therapy to Treat Multidrug-Resistant Bacterial Infections at a Single Center in the United States. Open Forum Infect Dis 7:ofaa389.

15. Mendes JJ, Leandro C, Corte-Real S, Barbosa R, Cavaco-Silva P, Melo-Cristino J, Gorski A, Garcia M. 2013. Wound healing potential of topical bacteriophage therapy on diabetic cutaneous wounds. Wound Repair Regen 21:595–603.

16. CDC. 2019. Antibiotic Resistance Threats in the United States, 2019. Atlanta, GA: U.S. Department of Health and Human Services.

17. Restrepo MI, Babu BL, Reyes LF, Chalmers JD, Soni NJ, Sibila O, Faverio P, Cilloniz C, Rodriguez-Cintron W, Aliberti S, Glimp. 2018. Burden and risk factors for Pseudomonas aeruginosa community-acquired pneumonia: a multinational point prevalence study of hospitalised patients. Eur Respir J 52.

18. Foundation CF. 2022. Cystic Fibrosis Foundation Patient Registry 2021 Annual Data Report. Bethesda, Maryland.

19. Kerem E, Reisman J, Corey M, Canny GJ, Levison H. 1992. Prediction of mortality in patients with cystic fibrosis. N Engl J Med 326:1187–91.

20. Strathdee SA, Hatfull GF, Mutalik VK, Schooley RT. 2023. Phage therapy: From biological mechanisms to future directions. Cell 186:17–31.

21. Sabourian P, Yazdani G, Ashraf SS, Frounchi M, Mashayekhan S, Kiani S, Kakkar A. 2020. Effect of Physico-Chemical Properties of Nanoparticles on Their Intracellular Uptake. Int J Mol Sci 21.

22. Li J, Mao H, Kawazoe N, Chen G. 2017. Insight into the interactions between nanoparticles and cells. Biomater Sci 5:173–189.

23. BK Chan GS, KE Kortright, M Modak, IM Ott, Y Sun, S Würstle, C Grun, B Kazmierczak, G Rajagopalan, Z Harris, CJ Britto, J Stewart, JS Talwalkar, C Appell, N Chaudary, SK Jagpal, R Jain, A Kanu, BS Quon, JM Reynolds, QA Mai, V Shabanova, PE Turner, JL Koff. 2023. Personalized Inhaled Bacteriophage Therapy Decreases Multidrug-Resistant Pseudomonas aeruginosa. medRxiv 10.1101/2023.01.23.22283996.

24. Nordstrom HR, Evans DR, Finney AG, Westbrook KJ, Zamora PF, Hofstaedter CE, Yassin MH, Pradhan A, Iovleva A, Ernst RK, Bomberger JM, Shields RK, Doi Y, Van Tyne D. 2022. Genomic characterization of lytic bacteriophages targeting genetically diverse Pseudomonas aeruginosa clinical isolates. iScience 25:104372.

25. Ehrhardt C, Collnot EM, Baldes C, Becker U, Laue M, Kim KJ, Lehr CM. 2006. Towards an in vitro model of cystic fibrosis small airway epithelium: characterisation of the human bronchial epithelial cell line CFBE41o. Cell Tissue Res 323:405–15.

26. Pezzulo AA, Tang XX, Hoegger MJ, Abou Alaiwa MH, Ramachandran S, Moninger TO, Karp PH, Wohlford-Lenane CL, Haagsman HP, van Eijk M, Banfi B, Horswill AR, Stoltz DA, McCray PB, Jr., Welsh MJ, Zabner J. 2012. Reduced airway surface pH impairs bacterial killing in the porcine cystic fibrosis lung. Nature 487:109–13.

27. Coakley RD, Grubb BR, Paradiso AM, Gatzy JT, Johnson LG, Kreda SM, O’Neal WK, Boucher RC. 2003. Abnormal surface liquid pH regulation by cultured cystic fibrosis bronchial epithelium. Proc Natl Acad Sci U S A 100:16083–8.

28. Stoltz DA, Meyerholz DK, Welsh MJ. 2015. Origins of cystic fibrosis lung disease. N Engl J Med 372:351–62.

29. Zajac M, Dreano E, Edwards A, Planelles G, Sermet-Gaudelus I. 2021. Airway Surface Liquid pH Regulation in Airway Epithelium Current Understandings and Gaps in Knowledge. Int J Mol Sci 22.

30. Katoh H, Fujita Y. 2012. Epithelial homeostasis: elimination by live cell extrusion. Curr Biol 22:R453–5.

31. Eisenhoffer GT, Loftus PD, Yoshigi M, Otsuna H, Chien CB, Morcos PA, Rosenblatt J. 2012. Crowding induces live cell extrusion to maintain homeostatic cell numbers in epithelia. Nature 484:546–9.

32. Conese M, Di Gioia S. 2021. Pathophysiology of Lung Disease and Wound Repair in Cystic Fibrosis. Pathophysiology 28:155–188.

33. Hornung V, Rothenfusser S, Britsch S, Krug A, Jahrsdorfer B, Giese T, Endres S, Hartmann G. 2002. Quantitative expression of toll-like receptor 1-10 mRNA in cellular subsets of human peripheral blood mononuclear cells and sensitivity to CpG oligodeoxynucleotides. J Immunol 168:4531–7.

34. Cao L, Chang H, Shi X, Peng C, He Y. 2016. Keratin mediates the recognition of apoptotic and necrotic cells through dendritic cell receptor DEC205/CD205. Proc Natl Acad Sci U S A 113:13438–13443.

35. Oganesyan V, Damschroder MM, Cook KE, Li Q, Gao C, Wu H, Dall’Acqua WF. 2014. Structural insights into neonatal Fc receptor-based recycling mechanisms. J Biol Chem 289:7812–24.

36. Stroh LJ, Stehle T. 2014. Glycan Engagement by Viruses: Receptor Switches and Specificity. Annu Rev Virol 1:285–306.

37. Caffrey M, Lavie A. 2021. pH-Dependent Mechanisms of Influenza Infection Mediated by Hemagglutinin. Front Mol Biosci 8:777095.

38. Morrison CB, Shaffer KM, Araba KC, Markovetz MR, Wykoff JA, Quinney NL, Hao S, Delion MF, Flen AL, Morton LC, Liao J, Hill DB, Drumm ML, O’Neal WK, Kesimer M, Gentzsch M, Ehre C. 2022. Treatment of cystic fibrosis airway cells with CFTR modulators reverses aberrant mucus properties via hydration. Eur Respir J 59.

39. Ludovico A, Moran O, Baroni D. 2022. Modulator Combination Improves In Vitro the Microrheological Properties of the Airway Surface Liquid of Cystic Fibrosis Airway Epithelia. Int J Mol Sci 23.

40. Abou Alaiwa MH, Launspach JL, Grogan B, Carter S, Zabner J, Stoltz DA, Singh PK, McKone EF, Welsh MJ. 2018. Ivacaftor-induced sweat chloride reductions correlate with increases in airway surface liquid pH in cystic fibrosis. JCI Insight 3.

41. Birket SE, Davis JM, Fernandez-Petty CM, Henderson AG, Oden AM, Tang L, Wen H, Hong J, Fu L, Chambers A, Fields A, Zhao G, Tearney GJ, Sorscher EJ, Rowe SM. 2020. Ivacaftor Reverses Airway Mucus Abnormalities in a Rat Model Harboring a Humanized G551D-CFTR. Am J Respir Crit Care Med 202:1271–1282.

42. Redrejo-Rodriguez M, Munoz-Espin D, Holguera I, Mencia M, Salas M. 2012. Functional eukaryotic nuclear localization signals are widespread in terminal proteins of bacteriophages. Proc Natl Acad Sci U S A 109:18482–7.

43. Zhang L, Sun L, Wei R, Gao Q, He T, Xu C, Liu X, Wang R. 2017. Intracellular Staphylococcus aureus Control by Virulent Bacteriophages within MAC-T Bovine Mammary Epithelial Cells. Antimicrob Agents Chemother 61.

44. Gorski A, Jonczyk-Matysiak E, Lusiak-Szelachowska M, Miedzybrodzki R, Weber-Dabrowska B, Borysowski J. 2018. Bacteriophages targeting intestinal epithelial cells: a potential novel form of immunotherapy. Cell Mol Life Sci 75:589–595.

45. Jurczak-Kurek A, Gasior T, Nejman-Falenczyk B, Bloch S, Dydecka A, Topka G, Necel A, Jakubowska-Deredas M, Narajczyk M, Richert M, Mieszkowska A, Wrobel B, Wegrzyn G, Wegrzyn A. 2016. Biodiversity of bacteriophages: morphological and biological properties of a large group of phages isolated from urban sewage. Sci Rep 6:34338.

46. Bach MS, de Vries CR, Khosravi A, Sweere JM, Popescu MC, Chen Q, Demirdjian S, Hargil A, Van Belleghem JD, Kaber G, Hajfathalian M, Burgener EB, Liu D, Tran QL, Dharmaraj T, Birukova M, Sunkari V, Balaji S, Ghosh N, Mathew-Steiner SS, El Masry MS, Keswani SG, Banaei N, Nedelec L, Sen CK, Chandra V, Secor PR, Suh GA, Bollyky PL. 2022. Filamentous bacteriophage delays healing of Pseudomonas-infected wounds. Cell Rep Med 3:100656.

47. Secor PR, Sweere JM, Michaels LA, Malkovskiy AV, Lazzareschi D, Katznelson E, Rajadas J, Birnbaum ME, Arrigoni A, Braun KR, Evanko SP, Stevens DA, Kaminsky W, Singh PK, Parks WC, Bollyky PL. 2015. Filamentous Bacteriophage Promote Biofilm Assembly and Function. Cell Host Microbe 18:549–59.

48. Burgener EB, Sweere JM, Bach MS, Secor PR, Haddock N, Jennings LK, Marvig RL, Johansen HK, Rossi E, Cao X, Tian L, Nedelec L, Molin S, Bollyky PL, Milla CE. 2019. Filamentous bacteriophages are associated with chronic Pseudomonas lung infections and antibiotic resistance in cystic fibrosis. Sci Transl Med 11.

49. Pinezich MR, Tamargo MA, Fleischer S, Reimer JA, Hudock MR, Hozain AE, Kaslow SR, Tipograf Y, Soni RK, Gavaudan OP, Guenthart BA, Marboe CC, Bacchetta M, O’Neill JD, Dorrello NV, Vunjak-Novakovic G. 2022. Pathological remodeling of distal lung matrix in end-stage cystic fibrosis patients. J Cyst Fibros 21:1027–1035.

50. McClure R, Massari P. 2014. TLR-Dependent Human Mucosal Epithelial Cell Responses to Microbial Pathogens. Front Immunol 5:386.

51. Li D, Wu M. 2021. Pattern recognition receptors in health and diseases. Signal Transduct Target Ther 6:291.

52. Shepardson KM, Schwarz B, Larson K, Morton RV, Avera J, McCoy K, Caffrey A, Harmsen A, Douglas T, Rynda-Apple A. 2017. Induction of Antiviral Immune Response through Recognition of the Repeating Subunit Pattern of Viral Capsids Is Toll-Like Receptor 2 Dependent. mBio 8.

53. Sartorius R, Trovato M, Manco R, D’Apice L, De Berardinis P. 2021. Exploiting viral sensing mediated by Toll-like receptors to design innovative vaccines. NPJ Vaccines 6:127.

54. Moussa EM, Panchal JP, Moorthy BS, Blum JS, Joubert MK, Narhi LO, Topp EM. 2016. Immunogenicity of Therapeutic Protein Aggregates. J Pharm Sci 105:417–430.

55. Sweere JM, Van Belleghem JD, Ishak H, Bach MS, Popescu M, Sunkari V, Kaber G, Manasherob R, Suh GA, Cao X, de Vries CR, Lam DN, Marshall PL, Birukova M, Katznelson E, Lazzareschi DV, Balaji S, Keswani SG, Hawn TR, Secor PR, Bollyky PL. 2019. Bacteriophage trigger antiviral immunity and prevent clearance of bacterial infection. Science 363.

56. Suda T, Hanawa T, Tanaka M, Tanji Y, Miyanaga K, Hasegawa-Ishii S, Shirato K, Kizaki T, Matsuda T. 2022. Modification of the immune response by bacteriophages alters methicillin-resistant Staphylococcus aureus infection. Sci Rep 12:15656.

57. Gorski A, Bollyky PL, Przybylski M, Borysowski J, Miedzybrodzki R, Jonczyk-Matysiak E, Weber-Dabrowska B. 2018. Perspectives of Phage Therapy in Non-bacterial Infections. Front Microbiol 9:3306.

58. Gogokhia L, Buhrke K, Bell R, Hoffman B, Brown DG, Hanke-Gogokhia C, Ajami NJ, Wong MC, Ghazaryan A, Valentine JF, Porter N, Martens E, O’Connell R, Jacob V, Scherl E, Crawford C, Stephens WZ, Casjens SR, Longman RS, Round JL. 2019. Expansion of Bacteriophages Is Linked to Aggravated Intestinal Inflammation and Colitis. Cell Host Microbe 25:285–299 e8.

59. Wang G, Nauseef WM. 2022. Neutrophil dysfunction in the pathogenesis of cystic fibrosis. Blood 139:2622–2631.

60. Abedon ST, Danis-Wlodarczyk KM, Wozniak DJ. 2021. Phage Cocktail Development for Bacteriophage Therapy: Toward Improving Spectrum of Activity Breadth and Depth. Pharmaceuticals (Basel) 14.

61. Pezzulo AA, Starner TD, Scheetz TE, Traver GL, Tilley AE, Harvey BG, Crystal RG, McCray PB, Jr., Zabner J. 2011. The air-liquid interface and use of primary cell cultures are important to recapitulate the transcriptional profile of in vivo airway epithelia. Am J Physiol Lung Cell Mol Physiol 300:L25–31.

62. Garcia-Castillo MD, Chinnapen DJ, Lencer WI. 2017. Membrane Transport across Polarized Epithelia. Cold Spring Harb Perspect Biol 9.

63. Barr JJ, Auro R, Furlan M, Whiteson KL, Erb ML, Pogliano J, Stotland A, Wolkowicz R, Cutting AS, Doran KS, Salamon P, Youle M, Rohwer F. 2013. Bacteriophage adhering to mucus provide a non-host-derived immunity. Proc Natl Acad Sci U S A 110:10771–6.

64. Carroll-Portillo A, Lin HC. 2021. Exploring Mucin as Adjunct to Phage Therapy. Microorganisms 9.

65. Lusiak-Szelachowska M, Miedzybrodzki R, Rogoz P, Weber-Dabrowska B, Zaczek M, Gorski A. 2022. Do Anti-Phage Antibodies Persist after Phage Therapy? A Preliminary Report. Antibiotics (Basel) 11.

66. Kazmierczak Z, Majewska J, Miernikiewicz P, Miedzybrodzki R, Nowak S, Harhala M, Lecion D, Keska W, Owczarek B, Ciekot J, Drab M, Kedzierski P, Mazurkiewicz-Kania M, Gorski A, Dabrowska K. 2021. Immune Response to Therapeutic Staphylococcal Bacteriophages in Mammals: Kinetics of Induction, Immunogenic Structural Proteins, Natural and Induced Antibodies. Front Immunol 12:639570.

67. Hodyra-Stefaniak K, Kazmierczak Z, Majewska J, Sillankorva S, Miernikiewicz P, Miedzybrodzki R, Gorski A, Azeredo J, Lavigne R, Lecion D, Nowak S, Harhala M, Wasko P, Owczarek B, Gembara K, Dabrowska K. 2020. Natural and Induced Antibodies Against Phages in Humans: Induction Kinetics and Immunogenicity for Structural Proteins of PB1-Related Phages. Phage (New Rochelle) 1:91–99.

68. Suh GA, Lodise TP, Tamma PD, Knisely JM, Alexander J, Aslam S, Barton KD, Bizzell E, Totten KMC, Campbell JL, Chan BK, Cunningham SA, Goodman KE, Greenwood-Quaintance KE, Harris AD, Hesse S, Maresso A, Nussenblatt V, Pride D, Rybak MJ, Sund Z, van Duin D, Van Tyne D, Patel R, Antibacterial Resistance Leadership G. 2022. Considerations for the Use of Phage Therapy in Clinical Practice. Antimicrob Agents Chemother 66:e0207121.

69. Van Belleghem JD, Clement F, Merabishvili M, Lavigne R, Vaneechoutte M. 2017. Pro- and anti-inflammatory responses of peripheral blood mononuclear cells induced by Staphylococcus aureus and Pseudomonas aeruginosa phages. Sci Rep 7:8004.

70. Bonilla N, Rojas MI, Netto Flores Cruz G, Hung SH, Rohwer F, Barr JJ. 2016. Phage on tap-a quick and efficient protocol for the preparation of bacteriophage laboratory stocks. PeerJ 4:e2261.

71. Mandrup OA, Lykkemark S, Kristensen P. 2017. Targeting of phage particles towards endothelial cells by antibodies selected through a multi-parameter selection strategy. Sci Rep 7:42230.

72. Marchesi JR, Sato T, Weightman AJ, Martin TA, Fry JC, Hiom SJ, Dymock D, Wade WG. 1998. Design and evaluation of useful bacterium-specific PCR primers that amplify genes coding for bacterial 16S rRNA. Appl Environ Microbiol 64:795–9.

73. Illumina. 2021. BCL Convert: a proprietary Illumina software for the conversion of BCL files to basecalls.

74. Bolger A, Giorgi F. 2014. Trimmomatic: A flexible read trimming tool for Illumina NGS data. Bioinformatics 30:2114–2120.

75. Seemann T. 2014. Prokka: rapid prokaryotic genome annotation. Bioinformatics 30:2068–9.

76. Meier-Kolthoff JP, Goker M. 2017. VICTOR: genome-based phylogeny and classification of prokaryotic viruses. Bioinformatics 33:3396–3404.

77. Meier-Kolthoff JP, Auch AF, Klenk HP, Goker M. 2013. Genome sequence-based species delimitation with confidence intervals and improved distance functions. BMC Bioinformatics 14:60.

78. Ye J, Coulouris G, Zaretskaya I, Cutcutache I, Rozen S, Madden TL. 2012. Primer-BLAST: a tool to design target-specific primers for polymerase chain reaction. BMC Bioinformatics 13:134.

79. Turner D, Shkoporov AN, Lood C, Millard AD, Dutilh BE, Alfenas-Zerbini P, van Zyl LJ, Aziz RK, Oksanen HM, Poranen MM, Kropinski AM, Barylski J, Brister JR, Chanisvili N, Edwards RA, Enault F, Gillis A, Knezevic P, Krupovic M, Kurtboke I, Kushkina A, Lavigne R, Lehman S, Lobocka M, Moraru C, Moreno Switt A, Morozova V, Nakavuma J, Reyes Munoz A, Rumnieks J, Sarkar BL, Sullivan MB, Uchiyama J, Wittmann J, Yigang T, Adriaenssens EM. 2023. Abolishment of morphology-based taxa and change to binomial species names: 2022 taxonomy update of the ICTV bacterial viruses subcommittee. Arch Virol 168:74.

80. Andrews S. 2010. FastQC: a quality control tool for high throughput sequence data. Babraham Bioinformatics, Babraham Institute, Cambridge, United Kingdom.

81. Andrews S, Krueger F, Segonds-Pichon A, Biggins L, Virk B, Dalle-Pezze P, Wingett S, Saadeh H, Ahlfors H. 2015. Trim Galore. Trim Galore.

82. Dobin A, Davis CA, Schlesinger F, Drenkow J, Zaleski C, Jha S, Batut P, Chaisson M, Gingeras TR. 2013. STAR: ultrafast universal RNA-seq aligner. Bioinformatics 29:15–21.

83. Liao Y, Smyth GK, Shi W. 2014. featureCounts: an efficient general purpose program for assigning sequence reads to genomic features. Bioinformatics 30:923–30.

84. Ewels P, Magnusson M, Lundin S, Kaller M. 2016. MultiQC: summarize analysis results for multiple tools and samples in a single report. Bioinformatics 32:3047–8.

85. Robinson MD, McCarthy DJ, Smyth GK. 2010. edgeR: a Bioconductor package for differential expression analysis of digital gene expression data. Bioinformatics 26:139–40.

86. Blighe K, Rana S, Lewis M. 2019. EnhancedVolcano: Publication-ready volcano plots with enhanced colouring and labeling. doi: 10.18129/B9.bioc.EnhancedVolcano.

87. Kramer A, Green J, Pollard J, Jr., Tugendreich S. 2014. Causal analysis approaches in Ingenuity Pathway Analysis. Bioinformatics 30:523–30.

88. Armbruster CR, Marshall CW, Garber AI, Melvin JA, Zemke AC, Moore J, Zamora PF, Li K, Fritz IL, Manko CD, Weaver ML, Gaston JR, Morris A, Methe B, DePas WH, Lee SE, Cooper VS, Bomberger JM. 2021. Adaptation and genomic erosion in fragmented Pseudomonas aeruginosa populations in the sinuses of people with cystic fibrosis. Cell Rep 37:109829.

